# BLOOM: Biologically Informed Flow Matching for Transcriptomic Virtual Embryo Modeling

**DOI:** 10.64898/2026.10.26.761895

**Authors:** Zhuo Liang, Shumin Li, Beier Li, Yuanhua Huang, Lequan Yu

## Abstract

Constructing a virtual embryo requires a generative representation of how heterogeneous cell populations change across developmental time. Here we introduce BLOOM, a biologically informed framework for conditional generation of developmental cell populations. BLOOM integrates temporal population context with developmental and regulatory priors through conditional flow matching to generate cell-expression distributions at queried stages in either temporal direction. Evaluation spans three mouse single-cell atlases covering prenatal development, visual-cortex maturation and multi-organ endothelial specialization. In comparisons with 22 time-series forecasting baselines, BLOOM ranks first for gene- and cell-expression correlations across all three datasets. Measured-stage evaluations support recovery of developmental expression patterns, population organization and marker-specific temporal relationships. Biological analyses connect these predictions to cell-state identities, gene programmes and regulatory associations. At unmeasured stages, interpolation and autoregressive history and future rollouts extend simulated expression dynamics between and beyond measured timepoints. Analyses of marker genes and temporal descriptors identify candidate differences in the timing of muscle-associated, neural and vascular-specialization programmes. Input-based gene perturbations further probe predicted responses of edited genes and associated molecular programmes across time. Together, BLOOM supports transcriptomic virtual embryo modeling by bridging temporal sampling gaps, exploring extended developmental trajectories and prioritizing gene-dependent hypotheses for experimental investigation.

## 1 Introduction

Development transforms an initially simple population of cells into diverse tissues through coordinated changes in gene expression, cell identity and intercellular organization. A predictive understanding of this process requires models that can connect observed developmental states to the populations expected at other ages and explore how changes in molecular inputs alter those predictions. This objective is part of the broader effort to build virtual cells, organs and organisms: computational representations that support queries about biological states and their responses to changing conditions [1, 2]. The Virtual Embryo Challenge brings this ambition into a shared evaluation setting, with tasks for temporal gene-expression distribution prediction, spatial–temporal prediction and mutant perturbation prediction [3]. Within this broader goal, forecasting developmental transcriptomic populations is a tractable component of a virtual embryo, providing a molecular substrate for analyses of changing cell states without requiring a complete simulation of tissue morphogenesis.

Time-resolved single-cell atlases provide an increasingly rich foundation for this task. Early whole-embryo atlases in zebrafish and *Xenopus*, together with mouse gastrulation and organogenesis atlases, established that developmental state transitions can be examined across experimentally recorded stages and multiple lineages [4–7]. More recent mouse resources extend temporal coverage across distinct biological scales: a prenatal atlas spans late gastrulation (E8) to birth (P0) with staging at approximately 2–6-hour intervals [8]; a cortex-focused atlas samples 35 scRNA-seq ages extending from embryonic development through postnatal maturation to adulthood [9]; and STED-EC follows endothelial development across 26 timepoints and eight organs, enabling analyses of how a shared cellular compartment acquires organ-specific expression states [10]. Their depth creates opportunities to learn from multiple observed stages. Nevertheless, temporal span and sampling intervals vary, and destructive sequencing yields independently sampled populations with changing cellular composition. The modeling problem is therefore to connect a history of population snapshots across developmental time while retaining the heterogeneity represented within each stage.

Computational approaches have progressively expanded what can be inferred from these snapshots. Pseudotime methods, including Monocle and diffusion pseudotime, organize asynchronous cells along transcriptional progressions and branching paths [11, 12]. Predicting expression at a specified chronological age requires additional information about how populations change between measured stages. Waddington-OT uses optimal transport to infer correspondences between time-indexed populations [13]. PRESCIENT learns stochastic developmental dynamics using expression and proliferation information, supporting trajectory simulation and expression perturbations [14]. TrajectoryNet, MIOFlow and scNODE further develop continuous generative dynamics for reconstructing or predicting temporal populations [15–17]. These advances establish developmental prediction as a distributional learning problem and motivate closer attention to how models use available temporal observations, biological structure and population variability to predict held-out stages.

Flow matching provides a framework for learning generative vector fields through regression along specified probability paths [18]. In single-cell analysis, simulation-free score and flow matching reconstructs stochastic dynamics between observed populations [19]. Multi-marginal flow matching extends this formulation across multiple sampling times and experimental conditions, using smooth paths to integrate information across temporal observations [20]. Joint velocity–growth flow matching additionally models changes in cell-state distributions together with population growth [21]. More recently, scFM combines optimal-transport supervision, bidirectional velocity fields and distribution-level alignment to improve temporal coherence during extended rollouts [22]. These developments establish a foundation for generating cellular populations across incompletely sampled time courses.

Biological knowledge can guide these models through constraints on transitions, structured architectures or molecular mechanisms. TrajectoryNet incorporates growth, data-manifold and RNA-velocity constraints when learning population dynamics [15]. For longitudinal spatial omics, ContextFlow uses tissue organization and ligand– receptor communication to guide optimal-transport couplings [23]. Navigo combines population-level flow matching with equations describing RNA transcription, splicing and degradation, grounding developmental vector fields in molecular kinetics [24]. In perturbation prediction, scDFM uses a co-expression graph estimated from training cells to structure gene attention [25]. Together, these approaches demonstrate several ways to connect expression generation with biological organization, from relationships among genes and neighbouring cells to the molecular processes underlying expression changes.

The representation of observed populations and temporal context is another important consideration. CellFlow conditions expression-distribution predictions on perturbation and experimental context, including developmental time in its developmental application [26]. TempoBridge explicitly supplies a pooled source-population summary alongside a requested transition descriptor, allowing transport to depend on population context [27]. CellPace represents temporal sequences with a stage- and gap-conditioned diffusion-forcing Transformer and sliding-window generation, supporting simulation, interpolation and future prediction [28]. These advances motivate bringing ordered population histories together with biological knowledge spanning cell hierarchies, developmental state transitions, tissue context and gene regulation. Integrating these resources within a common conditional representation provides a basis for developmental queries informed by both the measured expression series and the biological relationships underlying the system.

Here we introduce BLOOM (**B**idirectional **L**earning of **O**rgan **O**ntogeny with Flow **M**atching), a framework that treats developmental expression modeling as conditional time-series generation given an observed population window, a requested stage and supplied cell-state information. A shared cell encoder and stage-level pooling represent independently sampled cell sets, and a recurrent temporal encoder summarizes their ordered history. Inferred pseudo-trajectories supply cross-stage expression correspondences. Dedicated encoders and learned gates integrate developmental age, hierarchical cell identities, curated developmental relationships, anatomical context, state-associated gene programmes and weighted transcription-factor–target networks. The resulting representation conditions both a biologically informed Gaussian base and a flow-matching generator, placing biological knowledge at the initialization and generation stages of the model. This design brings cell-to-cell temporal correspondences and knowledge about cell states, tissues and genes into the same developmental prediction framework.

We evaluate BLOOM across the prenatal, visual-cortex and multi-organ endothelial atlases using population-, gene- and programme-level analyses. In comparisons with 22 time-series forecasting baselines [29–50], BLOOM ranks first for stage-averaged gene and cell correlations in all three datasets, alongside recognizable population structure and gene-specific expression patterns. The framework supports later-stage prediction from earlier observations and earlier-stage reconstruction from later observations. Interpolation queries intermediate states between measured stages, while recursive history and future rollouts extend simulated expression before the first and beyond the last observation. Input-based gene perturbations further probe predicted responses of edited genes and associated molecular programmes across time. Together, these capabilities support a transcriptomic virtual embryo that connects sparse developmental measurements, extends the analysis of temporal expression programmes and prioritizes gene-dependent hypotheses for experimental investigation.

## 2 Results

### 2.1 BLOOM models developmental expression dynamics with biological priors

We developed BLOOM to infer cell-expression populations at requested developmental stages from time-resolved single-cell RNA sequencing measurements, providing a transcriptomic framework for virtual embryo modeling (Fig. 1a). In this study, a virtual embryo is represented through populations of developmental transcriptomic states. Each measured stage comprises a set of cells described over a fixed gene panel, with the number and composition of sampled cells varying across stages. BLOOM formulates prediction as conditional generation of gene-expression profiles given an observed temporal window, the requested developmental stage and supplied cell-state information. This formulation accommodates prediction from earlier observations and reconstruction from later observations, interpolation between measured stages, and pre-measurement history and post-measurement future rollouts, extending developmental analysis into intervals and stages where experimental profiles are sparse or unavailable.

**Fig. 1.**
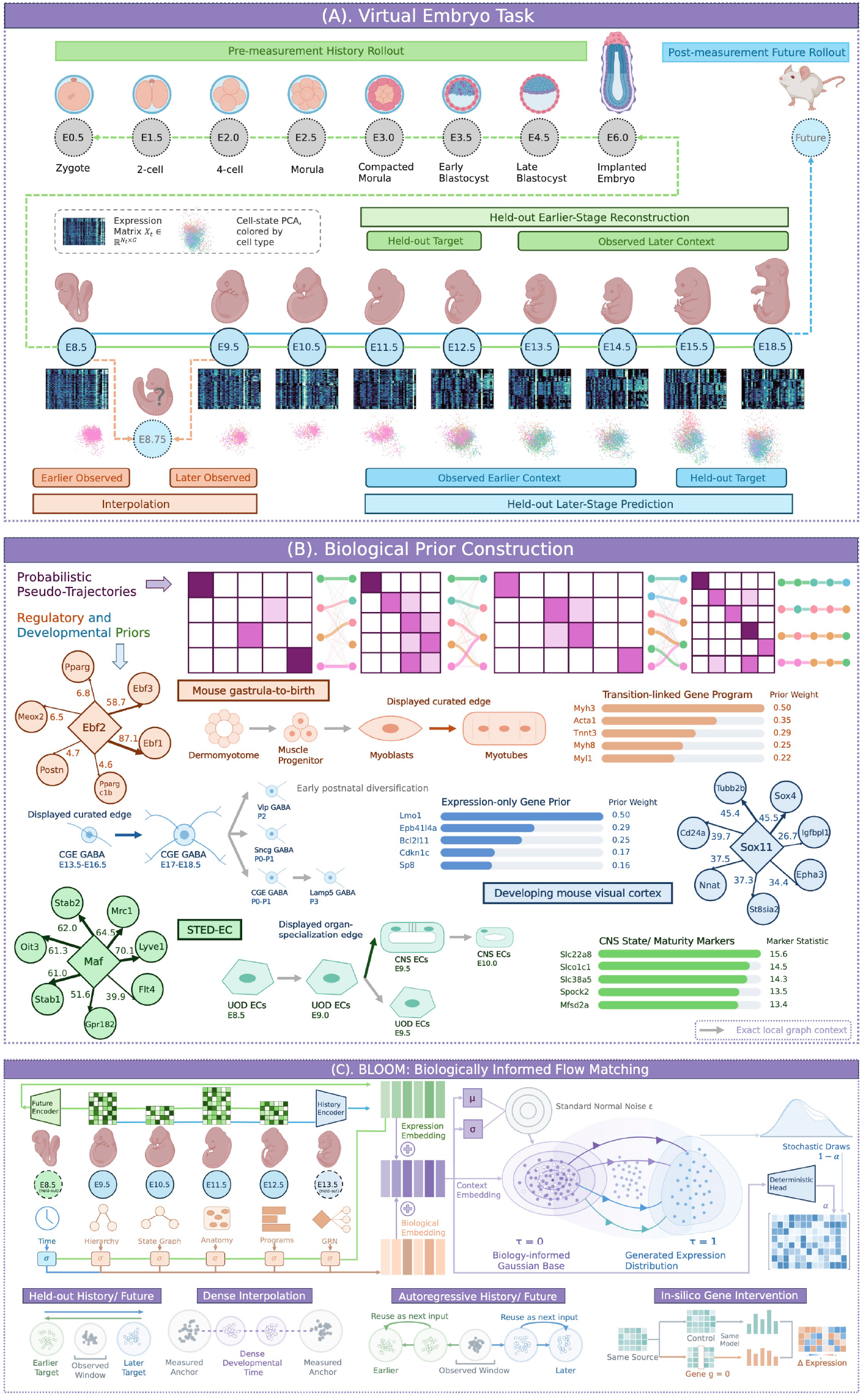
BLOOM integrates biological priors to model transcriptomic dynamics in a virtual embryo. **A,** Time-resolved single-cell measurements provide molecular snapshots of a changing developmental system. BLOOM connects these snapshots through five temporal queries: held-out later-stage prediction from earlier observations; held-out earlier-stage reconstruction from later observations; interpolation of unmeasured expression states between sampled stages; pre-measurement history rollout towards earlier development; and post-measurement future rollout beyond the sampled time range. **B,** Probabilistic cell-to-cell correspondences based on expression similarity across stages define pseudo-trajectories. For three datasets spanning mouse embryogenesis, visual cortex development and the endothelial atlas (STED-EC), published annotations and curated resources provide hierarchical cell identities, developmental relationships, anatomical context and state-associated gene programmes, complemented by inferred weighted transcription-factor– target networks. Representative networks and programmes illustrate the biological knowledge incorporated for these datasets. **C,** BLOOM integrates temporal population context, inferred cell correspondences and developmental and regulatory knowledge within a flow-matching generator. Applications include later-stage prediction, earlier-stage reconstruction, interpolation, history and future rollouts, and in silico gene perturbations. These capabilities complement sparse developmental experiments by extending temporal coverage and generating testable hypotheses about gene-dependent expression responses. GRN, gene regulatory network; *µ* and *σ*, Gaussian mean and standard deviation; *α*, weight of the deterministic prediction in the blended readout.

To connect independently sampled populations, BLOOM uses pseudo-trajectories as inferred temporal correspondences (Fig. 1b). Optimal transport provides a population-level basis for linking such cross-sectional measurements [13, 51]; the resulting links supply candidate predecessor or successor expression profiles without experimentally tracked cell histories. Complementary biological priors encode developmental age, hierarchical cell identities, curated developmental state relationships, anatomical context, state-associated gene programs and transcription-factor–target relationships. Weighted regulatory networks represent gene relationships inferred using single-cell network-inference frameworks [52, 53]. These resources enter BLOOM as conditioning features, complementing cell-to-cell correspondences with developmental, anatomical and gene-regulatory knowledge.

BLOOM first summarizes the observed populations with a shared cell encoder and permutation-invariant pooling within each stage. A recurrent temporal encoder (GRU) summarizes the stage representations, and its output is combined with cell-state information and pseudo-trajectory context. Dedicated feature encoders and learned gates further integrate biological priors into the conditional representation (Fig. 1c). Developmental graph features encode relationships among cell states, and regulatory-network features summarize expression through weighted transcription-factor–target connections. These inputs guide both initialization and expression generation. They specify the mean and standard deviation of a biologically informed Gaussian base. Samples from this base are then transformed towards the requested-stage expression distribution by a flow-matching velocity field [18]. A complementary deterministic head predicts endpoint expression from the same context, and this estimate can be combined with flow-generated profiles.

BLOOM supports inference in both developmental directions, predicting later-stage expression from earlier observations and reconstructing earlier-stage expression from later observations. Interpolation bridges gaps between measured stages to examine intermediate changes in cell-state expression and gene programs at finer temporal resolution. Recursive history and future rollouts extend simulated expression dynamics before the first and beyond the last measurement, enabling exploration of longer developmental time courses from limited observations. In-silico gene perturbations alter selected expression inputs to probe predicted responses of the edited gene and associated molecular programs across developmental time. Together, these capabilities complement sparse developmental experiments by filling temporal sampling gaps, extending simulated developmental trajectories and prioritizing gene-dependent responses for experimental investigation.

### 2.2 BLOOM preserves developmental population structure and cell-state identity

We next examined whether BLOOM preserves the organization of developmental cell-expression populations, extending assessment beyond individual expression values to population geometry, temporal progression and recognizable cell-state identity (Fig. 2). We considered three complementary developmental settings: the Mouse gastrula-to-birth atlas, which resolves extensive cellular diversification during prenatal organogenesis [8]; the Developing mouse visual cortex atlas, which documents continuing refinement of cortical cell identities through embryonic and postnatal development [9]; and STED-EC, which follows the acquisition of organ-associated endothelial identities during embryogenesis [10]. Together, these settings provide biologically distinct tests of whether conditional expression generation maintains structure across a broad embryonic landscape, a developing neural tissue and an organ-resolved vascular compartment.

**Fig. 2.**
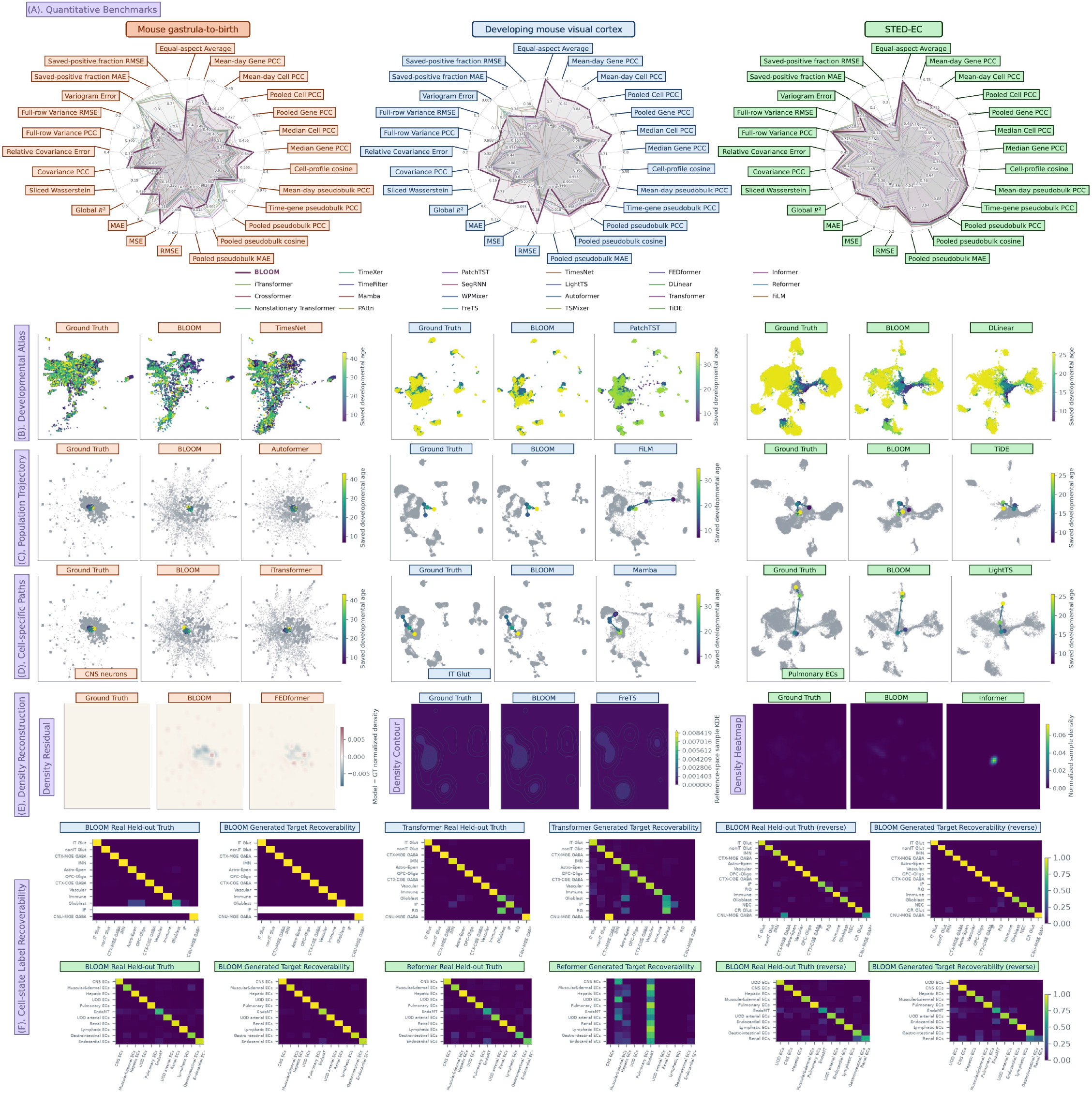
BLOOM reconstructs population geometry, stage-associated trajectories and cell-state signatures. **A,** Quantitative profiles for Mouse gastrula-to-birth, Developing mouse visual cortex and STED-EC. Spokes summarize complementary expression, correlation, pseudobulk and distributional readouts, with outward values indicating better performance. The equal-aspect average is an exploratory normalized aggregate across these aspects, with BLOOM ranking first in all three datasets. Throughout the paper, orange, blue and green boxes denote these three datasets, respectively. **B,** Developmental expression atlases. Observed and generated profiles are displayed in an observed-expression reference geometry and coloured by their saved developmental-stage coordinate. BLOOM retains recognizable stage-associated expression regions, whereas PatchTST underrepresents late-stage-coloured cortical profiles and DLinear alters the geometry of endothelial expression regions. **C,** Population centroid trajectories. Coloured points mark stage-specific expression centroids and arrows connect them in stage order. Grey points show the observed or generated cell population belonging to the corresponding subpanel. BLOOM more closely follows the observed centroid arrangement, whereas FiLM and TiDE show displaced cortical and endothelial stage trajectories, respectively. **D,** Cell-state centroid trajectories for CNS neurons in Mouse gastrula-to-birth, IT Glut in Developing mouse visual cortex and pulmonary endothelial cells in STED-EC. Centroid paths summarize expression geometry within the selected categories across developmental stages. BLOOM retains the broad observed progression, whereas Mamba [37] distorts the IT Glut path and LightTS [36] shortens and displaces the pulmonary endothelial path. **E,** Population density reconstruction. Mouse gastrula-to-birth uses a normalized-density residual (generated minus observed reference), Developing mouse visual cortex uses density contours, and STED-EC uses a normalized-density heatmap. The observed residual is zero by construction. Colour scales have different meanings between these three display types, and density is normalized transcriptomic occupancy. BLOOM retains distributed endothelial occupancy, whereas Informer concentrates profiles in a restricted region. **F,** Real-cell classifier validation and generated target-label recoverability for forward and reverse diagnostics. Within each dataset row, the six columns show BLOOM real held-out expression, BLOOM generated expression, baseline real held-out expression, baseline generated expression, BLOOM reverse real held-out expression and BLOOM reverse generated expression. Rows denote supplied evaluation labels and columns denote classifier assignments. Colours show fractions normalized by the full original row support; white rows have no evaluable support. Class order can differ between forward and reverse exports. Real held-out expression calibrates classifier separability; generated-expression agreement measures conditional target-label recovery. Reverse diagnostics are evaluated with observed expression references. BLOOM retains pronounced diagonal recovery in both directions, whereas the forward Transformer and Reformer examples show greater cross-state assignment. Different representative baselines are displayed across dataset-specific views and tasks to illustrate the wider comparison. CNS, central nervous system; IT Glut, intratelencephalic glutamatergic neurons; EC, endothelial cell.

The quantitative profiles summarize complementary aspects of predictive agreement across BLOOM and 22 time-series baselines (Fig. 2a). BLOOM ranked first for stage-averaged gene Pearson correlation in all three datasets, with values of 0.532, 0.634 and 0.488 for the prenatal, visual-cortex and endothelial settings, respectively. Stage-averaged cell correlations were 0.427, 0.864 and 0.814, also ranking first in all three comparisons. BLOOM additionally achieved the highest global *R*^2^ (0.466, 0.762 and 0.673) and covariance-edge Pearson correlation (0.958, 0.976 and 0.921), supporting agreement in expression values and gene co-variation. In the visual-cortex and endothelial settings, BLOOM also achieved the lowest relative covariance error (0.257 and 0.410). The exploratory equal-aspect average ranked BLOOM first in all three datasets (0.730, 0.993 and 0.955). We further examined population organization through atlas, centroid, density and classifier views.

In the developmental expression atlases, BLOOM-generated profiles populate multiple regions of the observed transcriptomic reference geometry, with broadly recognizable distributions of the saved developmental-stage coordinate (Fig. 2b). The visual-cortex panels retain a prominent central population together with separated peripheral regions, while the endothelial panels retain several distinct expression regions rather than a single undifferentiated cloud. By comparison, PatchTST [40] underrepresents late-stage-coloured visual-cortex profiles, while DLinear [31] alters the separation and extent of endothelial expression regions. The whole-embryo setting additionally exposes differences in local dispersion, making it useful for assessing heterogeneity across a broad collection of developmental states. This population-level interpretation is consistent with treating developmental single-cell measurements as time-indexed expression distributions [13].

Population centroid trajectories summarize how these distributions change across the sampled stages (Fig. 2c). The visual-cortex BLOOM output retains a compact sequence of stage-associated centroids, and the endothelial output preserves a structured displacement across its displayed stages. The whole-embryo centroid sequence remains concentrated within the central expression region, despite appreciable variation in the surrounding cell clouds. This distinction is biologically informative: a stable pooled centroid can coexist with substantial heterogeneity or changing contributions from different states. The centroid views therefore complement the atlas views by displaying temporal organization, while the surrounding distributions retain information that an average position cannot capture.

We then examined CNS neurons, intratelencephalic glutamatergic neurons (IT Glut) and pulmonary endothelial cells to assess temporal structure within defined biological compartments (Fig. 2d). In the prenatal atlas, CNS neurons form a meaningful neural compartment within a much broader landscape of organogenesis [8]. Their compact centroid sequence provides a within-compartment check alongside the pooled embryonic population. In the visual cortex, the IT Glut class contains finer neuronal subclasses whose transcriptomic identities continue to develop over time [9]. BLOOM retains a stage-ordered progression within this class, supporting the preservation of temporal expression structure within a broad neuronal identity. This interpretation allows both maturation-related expression changes and shifts in the mixture of sub-classes within the class. Pulmonary endothelial cells provide a complementary vascular example: the STED-EC study identifies a conserved pulmonary endothelial transcriptional transition during organ-associated differentiation [10], and independent work establishes the developmental specialization of pulmonary capillary endothelial cells [54]. The BLOOM pulmonary centroid path retains the broad movement between expression regions seen in the observed panel. This supports preservation of organ-associated temporal structure at the displayed class resolution, without assigning the trajectory to a particular capillary subtype or inferring vascular anatomy.

Density views provide a further assessment of how generated expression is distributed within the reference space (Fig. 2e). The whole-embryo residual map localizes regions where generated normalized density is above or below the displayed observed reference, making remaining discrepancies visible even when centroids occupy similar locations. The visual-cortex contours retain several recognizable density regions, and the STED-EC BLOOM heatmap represents a dispersed population with multiple occupied regions. By comparison, the displayed Informer [35] output has a conspicuous localized density concentration. These observations illustrate why population fidelity requires consideration of local concentration as well as average position.

Expression-based classification supplied a complementary readout of whether generated cells retain the transcriptional signatures associated with their requested states (Fig. 2f). Classifying cells from their expression profiles is an established way to assess recognizable cell identities [55]. We interpret generated target-label recovery alongside classifier performance on real held-out expression. In both the visual-cortex and endothelial forward panels, BLOOM-generated profiles show a pronounced diagonal pattern across the supported states. The displayed Transformer [48] and Reformer [41] outputs show more off-diagonal assignments, with particularly concentrated assignments in the endothelial Reformer panel. Thus, the illustrated BLOOM outputs retain class-discriminating expression structure across multiple states. White rows identify states without evaluable row support and are excluded from this interpretation.

State signatures also remain recognizable in the reverse diagnostics (Fig. 2f). The reverse readouts extend the assessment of BLOOM beyond forward prediction: expression generated from later context retains recognizable signatures of the requested states. This supports a learned conditional transcriptomic correspondence across developmental time in both directions. The compared baselines were evaluated in their earlier-to-later forecasting configurations, whereas BLOOM explicitly incorporates a later-to-earlier reconstruction pathway. Identity recovery establishes a biologically meaningful component of reverse reconstruction; age-resolved reverse expression agreement is the complementary test of whether an output also matches the requested earlier developmental stage. Across these views, BLOOM supports the generation of structured developmental expression populations with recognizable cell-state identity, providing a population-level basis for the gene-level analyses that follow.

### 2.3 BLOOM recovers gene-level expression structure and molecular heterogeneity

Having examined population organization and cell-state identity, we next assessed whether BLOOM preserves the gene-level expression structure underlying these developmental populations (Fig. 3). Reference-based gene screening, stage-resolved distributions, expression overlays in transcriptomic space and stage-wise expression summaries provide complementary views of predictive fidelity. Together, they examine whether reconstructed populations retain both the distribution of individual transcripts and their changing representation across developmental stages.

**Fig. 3.**
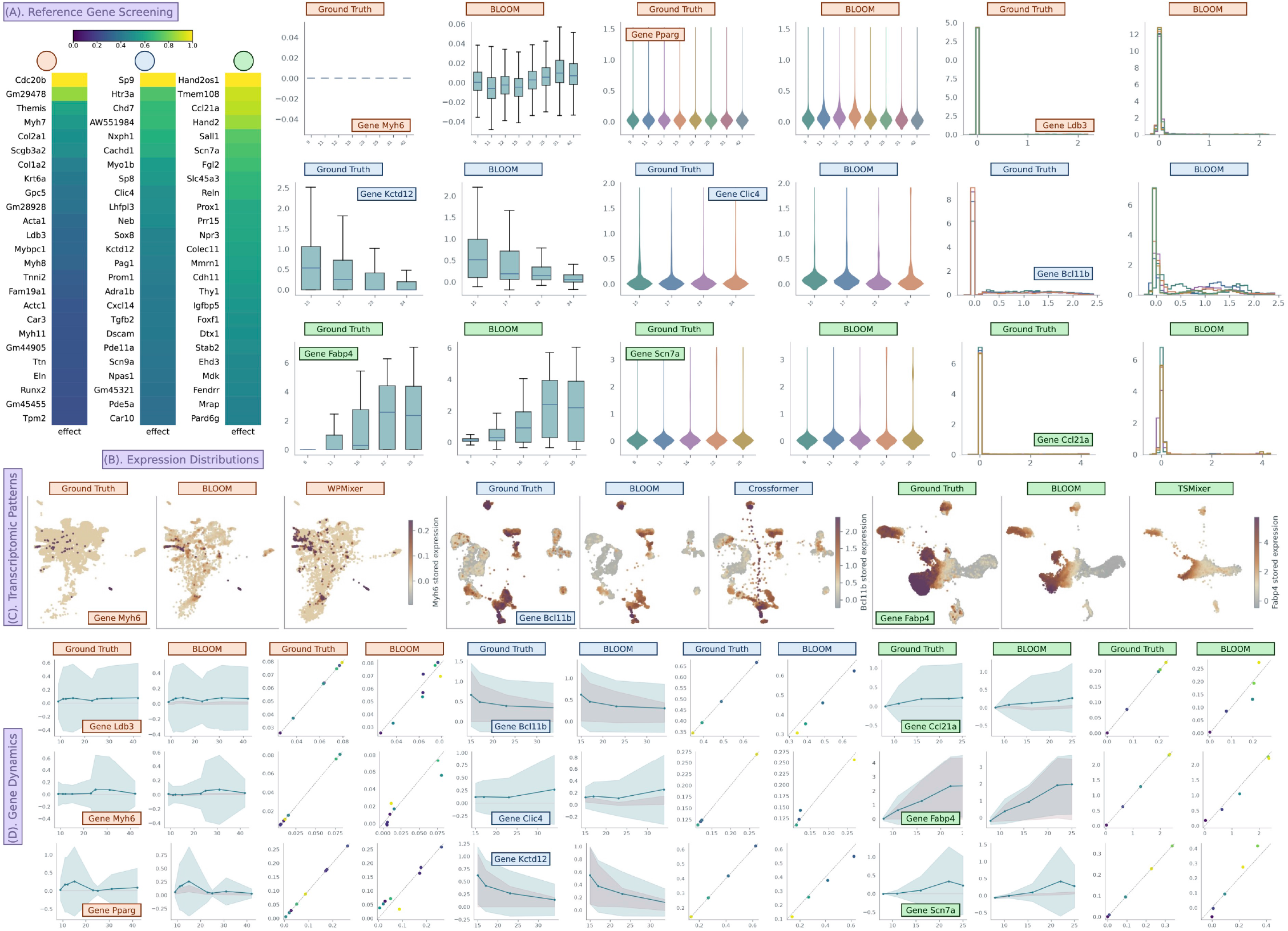
BLOOM reconstructs biologically informative gene-expression distributions, transcriptomic patterns and developmental trends. Orange, blue and green identify the Mouse gastrula-to-birth atlas, Developing mouse visual cortex and STED-EC, respectively. **A,** Reference gene screening using ground-truth expression. Each column shows the leading 25 genes ranked within the gene set curated from the corresponding dataset publication. The displayed genes are prioritized by cell-state-associated variation across sampled stages. Effect summarizes differences between observed group means relative to within-group expression variation. **B,** Stage-resolved expression distributions. Rows show the prenatal, cortical and endothelial datasets. From left to right, prenatal examples are *Myh6* boxplots, *Pparg* violins and *Ldb3* histograms; cortical examples are *Kctd12* boxplots, *Clic4* violins and *Bcl11b* histograms; endothelial examples are *Fabp4* boxplots, *Scn7a* violins and *Ccl21a* histograms. Each pair compares observed expression with BLOOM. Histograms and violin shapes are normalized within each displayed stage and expression range. Boxplots show sample medians and interquartile ranges, with 1.5-IQR (interquartile range) whiskers and outliers omitted. **C,** Gene-expression patterns in transcriptomic space. Observed-reference UMAP displays show *Myh6* in the prenatal dataset, *Bcl11b* in the cortical dataset and *Fabp4* in the endothelial dataset. Within each dataset, columns show observed expression, BLOOM and a representative baseline: WPMixer, Crossformer and TSMixer, respectively. Colour indicates gene expression on a shared display scale within each dataset comparison. Coordinates describe transcriptomic similarity. BLOOM retains recognizable gene-enriched regions across the three datasets. **D,** Developmental expression dynamics and stage-matched mean agreement. Rows within each dataset group show *Ldb3*, *Myh6* and *Pparg*; *Bcl11b*, *Clic4* and *Kctd12* ; and *Ccl21a*, *Fabp4* and *Scn7a*, respectively. Lines show full-population stage means; cyan bands denote mean ± cell standard deviation and pink bands denote the plotted sample’s interquartile range. Mean-comparison scatters use observed stage mean on the horizontal axis and the displayed source’s stage mean on the vertical axis. The diagonal denotes equality, and point colours encode saved stage index. Observed-versus-observed scatters lie on the diagonal by construction.

Reference gene screening used ground-truth expression to rank genes by biologically structured variation across the sampled developmental populations (Fig. 3a). The displayed columns show the leading 25 genes ranked within the gene sets curated from the corresponding dataset publications [8–10]. These genes were prioritized by cell-state-associated differences evaluated across developmental stages, providing reference sets of developmental indicator genes for subsequent visualization.

To complement the reference screening, we selected representative marker genes with established biological roles in each dataset. In the prenatal atlas, *Myh6*, *Ldb3* and *Pparg* span contractile and adipogenic-associated biology. *Myh6* encodes the alpha-cardiac myosin heavy chain; genetic ablation in mice establishes its importance for cardiac development and performance [56]. *Ldb3*, also known as *Cypher*, encodes a striated-muscle Z-line-associated PDZ-LIM protein whose loss disrupts Z-line integrity during muscle activity [57]. It therefore provides a readout of cardiac and skeletal-muscle-associated expression. *Pparg* provides a distinct regulatory example: genetic experiments establish PPAR*γ* as necessary for adipose differentiation [58]. Mouse knockout studies further establish its requirement for trophoblast differentiation and placental vascularization during embryogenesis [59]. These molecular roles make the three genes complementary tests of expression reconstruction.

The visual-cortex examples similarly cover different aspects of neuronal biology. *Bcl11b* encodes CTIP2, a transcription factor with experimentally demonstrated roles in neuronal projection development [60]. *Kctd12* encodes an auxiliary GABA_B_-receptor protein that modulates receptor signalling kinetics [61], providing a receptor-associated expression example. *Clic4* contributes an additional developmentally modulated readout: an independent knock-in reporter study documents fetal-brain expression alongside broad expression in other tissues [62]. Thus, the cortical examples connect developmental transcriptional regulation with receptor-associated and broader cellular expression programmes.

Within STED-EC, *Fabp4*, *Scn7a* and *Ccl21a* sample complementary molecular features of endothelial populations. *Fabp4* has experimentally established vascular expression and VEGF responsiveness, with a contribution to endothelial proliferation in cultured cells [63]. *Scn7a* supplies an ion-channel-associated specialization readout; its enrichment in renal glomerular endothelial cells has been documented by independent vascular expression profiling [64], supporting its relevance to endothelial heterogeneity. *Ccl21a* encodes the murine CCL21-Ser chemokine [65]. CCL21 presented on lymphatic endothelium contributes to dendritic-cell docking and migration into lymphatic vessels [66]. Together, these genes establish a biological context for assessing the reconstruction of endothelial specialization and signalling-associated transcripts.

Stage-resolved distributions show how this biological signal is represented beyond a population mean (Fig. 3b). The prenatal *Myh6* sample boxes collapse near zero in observed cells, whereas BLOOM produces a small nonzero spread. In the visual cortex, the *Kctd12* boxplots retain the broad decrease in expression across the displayed stages. In STED-EC, *Fabp4* distributions shift towards higher expression at later displayed stages in both observed and generated cells. The *Clic4* violin plots retain a concentration at low expression with extended positive tails, while *Scn7a* retains a low-expression body with a positive tail. The *Ldb3* and *Ccl21a* histograms preserve prominent near-zero components, illustrating the importance of reconstructing sparse expression as well as positive values. The *Bcl11b* histograms retain a broader positive-expression component, with differences in the near-zero peak. BLOOM thus captures gene-specific distributional patterns, including prominent near-zero components, positive-expression tails and shifts in expression level across stages.

Expression overlays then examine where individual genes are represented within the shared observed-expression UMAP (Fig. 3c). For prenatal *Myh6*, BLOOM retains a sparse, focal expression pattern in regions broadly corresponding to the observed reference, whereas WPMixer [49] distributes stronger signal across broader regions of the embedding. In the visual cortex, *Bcl11b* expression is enriched in separated regions, and BLOOM retains recognizable expression-rich compartments. The displayed Crossformer [30] reconstruction redistributes both points and expression intensity across the embedding. In STED-EC, BLOOM retains a substantial *Fabp4* - expressing region and several expression-rich peripheral compartments, whereas the displayed TSMixer [43] result shows a more diffuse pattern. These views support preservation of expression-associated organization within transcriptomic space.

Developmental timecourses provide another view of gene-level temporal structure (Fig. 3d). The prenatal *Pparg* profile retains an early local maximum followed by lower values; *Ldb3* and *Myh6* show comparatively modest mean variation over their displayed stages. BLOOM retains the broad decline in the cortical *Bcl11b* and *Kctd12* stage means, together with the more modest changes in *Clic4*. In STED-EC, *Fabp4* means rise towards later stages in both observed and generated populations, while *Ccl21a* and *Scn7a* show smaller-amplitude changes. These differences demonstrate why a useful temporal reconstruction must accommodate decreasing, increasing and non-monotonic expression patterns. The accompanying stage-mean comparisons place predicted and observed means against the equality line, making amplitude agreement and residual offsets visible.

The distributions and timecourses also summarize different aspects of the evaluated populations. Sample quantiles can remain at zero for sparse genes even when a full-population mean is positive; mean agreement therefore does not guarantee agreement in the fraction of expressing cells or in the positive tail. Likewise, a pooled stage mean can reflect both expression changes within cell states and changing contributions from different states. Considered together, the selected distribution, embedding and temporal views show that BLOOM reconstructs biologically meaningful gene-associated expression structure across distinct developmental systems, while retaining visible gene- and stage-specific discrepancies for further refinement.

### 2.4 BLOOM preserves cellular marker signatures and programme-associated expression

Beyond recovering individual gene distributions, a virtual embryo should retain the molecular combinations that distinguish cell populations and their developmental programmes. We therefore examined whether the gene-level agreement described above extends to coherent state-associated organization (Fig. 4). Linking marker expression to population composition and programme-associated patterns allows reconstructed cells to be interpreted in the biological contexts of the reference atlases.

**Fig. 4.**
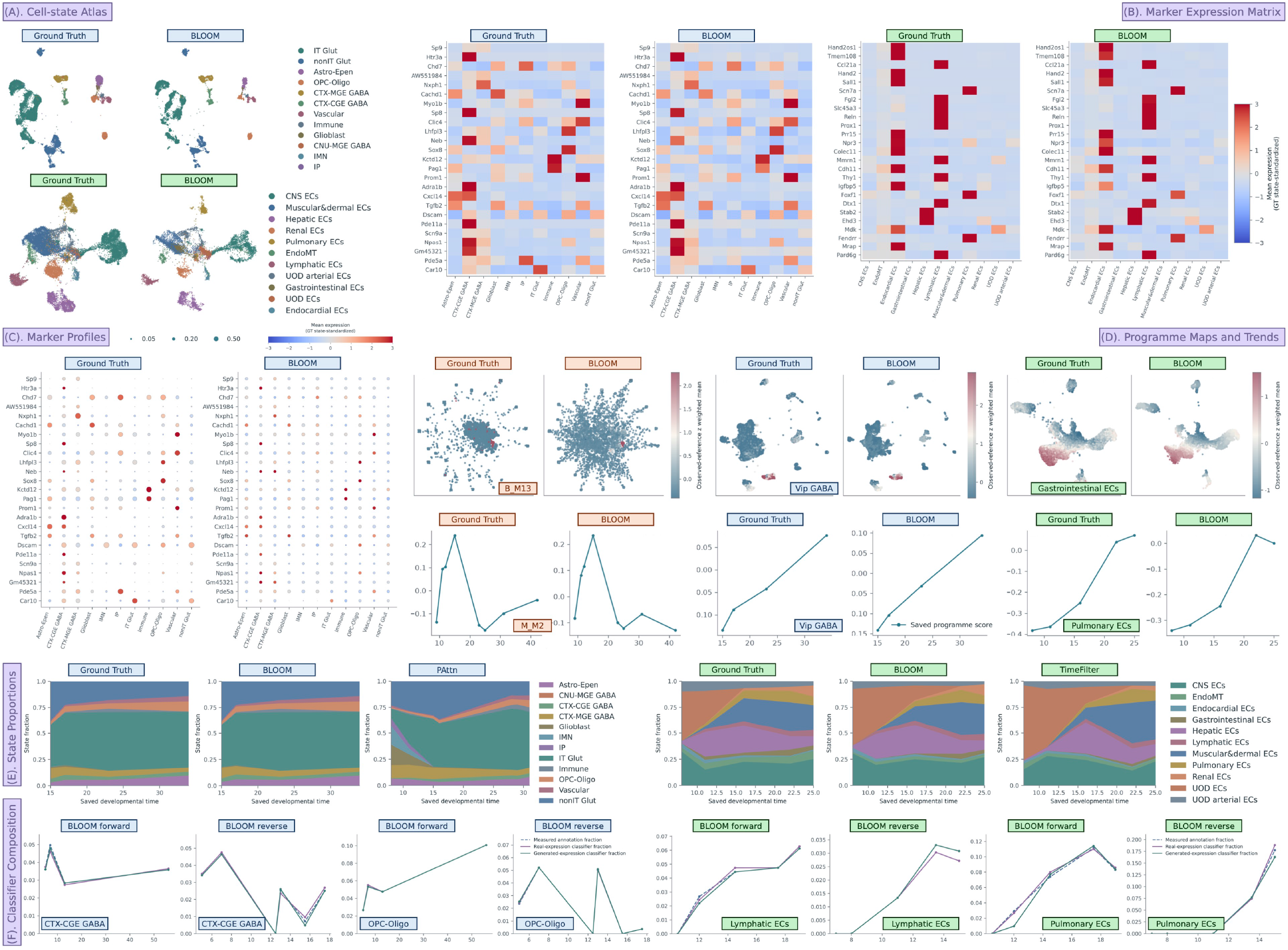
BLOOM retains state-associated marker patterns, population composition and localized developmental programme readouts. Orange, blue and green identify Mouse gastrula-to-birth, Developing mouse visual cortex and STED-EC, respectively. **A,** Reconstruction of annotated cell populations in transcriptomic reference space. Rows show visual cortex and STED-EC. Each row compares observed profiles with BLOOM in the observed-reference coordinate system. Colours denote reference-associated cell-state assignments. Generated state assignments are derived using the reference expression representation. **B,** State-resolved reconstruction of marker-expression patterns. The matrices compare selected top-25 marker means for visual cortex and STED-EC between observed populations and BLOOM. For each gene, colour is the cell-state mean centred on the equal-weight observed across-state mean and divided by the population standard deviation of those observed state means. All matrices use display bounds [−3, 3], with clipping only for colour. Rows denote genes and columns denote the supplied cell-state categories. **C,** Joint comparison of marker-expression intensity and expressing-cell fractions. Visual-cortex dotplots compare observed profiles with BLOOM using the same 25 genes and colour transform as **b**. Circle areas represent relative display fractions of stored expression greater than zero. **D,** Localization and temporal progression of selected developmental programme readouts. Upper maps show prenatal B M13, cortical VIP-associated score and gastrointestinal endothelial programme. Lower curves show prenatal M M2, cortical VIP-associated score and pulmonary endothelial programme. Scores are weighted means over represented programme genes, standardized to the observed expression reference. **E,** Stage-resolved reconstruction of annotated population composition. The visual-cortex and STED-EC panels compare observed proportions with BLOOM and representative PAttn or TimeFilter outputs, respectively. Fractions are reference-associated state occupancies and sum to one within each reported stage. **F,** Expression-based recovery of cell-class fractions in forward and reverse inference. Diagnostics compare forward and reverse inference for CTX-CGE GABA and OPC-Oligo in visual cortex, and lymphatic and pulmonary cells in STED-EC. The three traces represent measured annotation fractions, real-expression classifier fractions and generated-expression classifier fractions. They assess classifiability and composition within the supplied conditional evaluation. GT, ground truth; CGE/MGE, caudal/medial ganglionic eminence; IT Glut, intratelencephalic glutamatergic population; OPC-Oligo, oligodendrocyte precursor and oligodendrocyte category; EC, endothelial cell; EndoMT, supplied endothelial-to-mesenchymal transition annotation.

The cell-state atlases retain recognizable population organization in the observed-expression reference geometry (Fig. 4a). In the cortical atlas, BLOOM preserves the prominent intratelencephalic glutamatergic compartment together with smaller inhibitory, oligodendroglial and other annotated populations. In STED-EC, the reconstruction retains separated and adjoining compartments associated with distinct endothelial categories, including lymphatic, pulmonary, gastrointestinal and endocardial cells. These views establish a useful population-level context for evaluating molecular identity.

The marker matrices and cortical dotplots connect this organization to biologically informative expression patterns (Fig. 4b,c). In visual cortex, the pronounced *Htr3a* and *Sp8* enrichment in the CTX-CGE GABA category is retained by BLOOM. *Htr3a* has been experimentally characterized as an early and persistent marker of cortical interneuron subpopulations enriched for caudal ganglionic eminence origin [67]. Conditional genetic studies also establish contributions of *Sp8* and *Sp9* to the development and migration of CGE-derived interneurons [68]. The displayed *Sp9* signal is strongest in the CTX-MGE GABA category, illustrating the importance of interpreting combinations of markers within the actual reference rather than assigning an exclusive lineage from one transcript. *Cxcl14* adds a complementary developmental readout: its involvement in layer-I interneuron differentiation has been demonstrated in developing mouse somatosensory cortex [69], and its enriched cortical expression pattern remains recognizable in the reconstruction. The retention of *Sox8* enrichment in OPC-Oligo provides a distinct glial counterpart, consistent with the experimentally established oligodendroglial role of *Sox8* in the developing mouse spinal cord [70]. Together, these examples show preservation of molecular distinctions within both neuronal and non-neuronal compartments.

The endothelial marker panel provides particularly clear links between reconstructed expression and vascular specialization. BLOOM retains lymphatic enrichment of *Prox1* and *Ccl21a*, pulmonary enrichment of *Foxf1* and *Scn7a*, and endocardial enrichment of *Hand2* (Fig. 4b). *Prox1* is required for murine lymphatic development [71], whereas lymphatic endothelial CCL21 contributes to dendritic-cell passage across lymphatic endothelia [72]. Endothelial *Foxf1* supports embryonic vascular formation and VEGF signalling [73], and independent comparative endothelial profiling identifies pulmonary enrichment of *Scn7a* [74]. *Hand2* has an experimentally established role in atrioventricular cushion and cardiac valve development [75]. These complementary signatures make the panel a test of organ- and state-associated molecular structure, extending beyond a generic endothelial label.

The localized programme maps and representative curves resolve where and when selected developmental expression readouts occur (Fig. 4d). The prenatal B M13 and M M2 annotations correspond to Kupffer-cell and adipocyte-progenitor contexts in the supplied author graph. Tissue-resident macrophage specialization during organogenesis and adipogenic differentiation have distinct experimental foundations [76, 77], providing biological context for these expression summaries. The cortical VIP-associated score remains concentrated in a restricted transcriptomic compartment, and the gastrointestinal endothelial score retains a recognizable enriched region. The temporal examples illustrate complementary patterns: the prenatal M M2 readout rises to its maximum at saved stage 15 and subsequently falls, the cortical VIP-associated score increases across its four displayed stages, and the pulmonary endothelial readout shows a broad increase from earlier to later saved stages. BLOOM retains these principal shapes, preserving programme-associated expression changes that accompany cellular specialization across development.

State proportions then examine how reconstructed populations are represented across the saved developmental stages (Fig. 4e). In cortex, BLOOM retains the dominant IT Glut contribution and the broad arrangement of the less abundant categories. In STED-EC, it reproduces substantial changes in the relative contributions of the endothelial classes rather than maintaining a fixed composition across stages. The displayed PAttn [39] and TimeFilter [45] examples provide additional descriptive context, but their saved reporting grids or bound reference populations differ from those of the displayed BLOOM comparisons. The paired GT–BLOOM views therefore support the clearest interpretation of composition agreement within their native contexts.

Classifier-assigned fractions provide a complementary check that the supplied class composition remains recognizable from reconstructed expression (Fig. 4f). The cortical examples contrast CTX-CGE GABA with OPC-Oligo, bringing together an interneuron-associated category and a broad oligodendroglial category. The latter encompasses a developmental continuum rather than a single terminal state [78]. The endothelial examples contrast lymphatic and pulmonary categories whose molecular relevance is independently supported by the marker panel. In both forward and reverse diagnostics, the generated-expression classifier traces closely follow the corresponding real-expression traces across the displayed windows. Across the saved stages, their mean absolute fraction differences are below 0.19 percentage points for the selected cortical categories and below 0.57 percentage points for the selected endothelial categories.

Taken together, the cell-state, marker, composition and programme-associated views support BLOOM’s ability to retain biologically interpretable expression organization across distinct developmental systems. The strongest evidence comes from recognizable state-associated marker combinations and their accompanying population and temporal structure. Programme coverage and the conditional classifier design define the scope of that conclusion, while providing a basis for broader programme-level evaluation across more extensive gene panels.

### 2.5 BLOOM preserves temporal expression structure and biologically interpretable regulatory associations

We next examined whether BLOOM retains the temporal organization of gene expression together with the gene–gene associations and supplied regulatory modules that give those dynamics a biological context (Fig. 5). The analysis progresses from adjacent-stage expression ranks and gene-specific timing to coexpression, temporal mean-expression changes, candidate regulatory associations and multi-gene module scores. Across the mouse prenatal, developing visual-cortex and embryonic endothelial datasets [8–10], these complementary views assess expression organization at several resolutions.

**Fig. 5.**
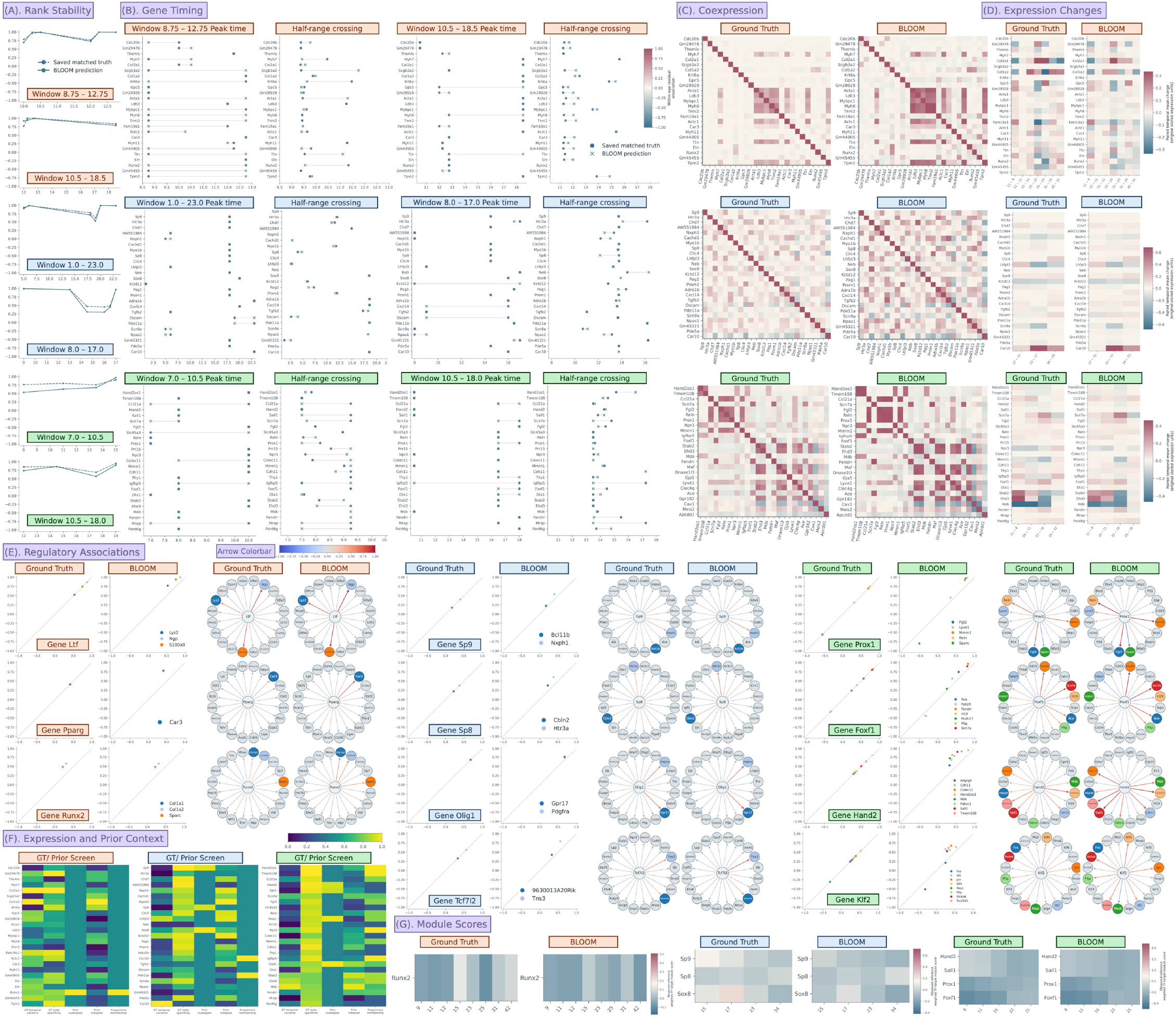
BLOOM preserves temporal expression structure and biologically interpretable regulatory associations. Orange, blue and green identify Mouse gastrula-to-birth, Developing mouse visual cortex and STED-EC, respectively. Reporting-time coordinates in **a,b** are supplied actual-time diagnostics; coordinates in **d,g** are saved stage indices. **A,** Persistence and reorganization of developmental gene-expression ranks. Spearman correlations compare gene-wise mean-expression ranks at consecutive sampled times, separately within observed and generated profiles. Points mark each interval’s right-hand time, and lines join successive comparisons. **B,** Gene-specific timing of expression peaks and transitions. The windows match **a**, with 25 selected genes per window. Peak time is the sampled-grid maximum. Half-range crossing is the first crossing of the profile midrange in its net direction, interpolated between observations where applicable. Points and crosses denote GT and BLOOM; segments pair the same gene. Crossings remain undefined when a profile starts beyond the threshold, lacks a net direction or does not cross within the sampled window. **C,** Coordination of gene expression within developmental stages. Matrices compare observed and generated Pearson correlations after within-age/stage centring of sampled expression. The common colour limits are [−1, 1]. **D,** Direction and magnitude of stage-to-stage expression changes. Adjacent-stage differences in weighted mean expression are shown for 25 selected temporal indicator genes. Each dataset has 20,000 matched cell/time records with equal supplied weights. Columns show target stage minus previous stage in stored expression units. GT and BLOOM share a colour scale within each dataset, with display bounds of approximately ±0.333, ±0.681 and ±0.465, respectively; clipping affects colour only. **E,** Reconstruction of development-associated gene–gene associations. Eleven examples pair GT/BLOOM scatterplots with circular prior networks. Scatter *x* is the observed-reference within-stage residual correlation and *y* is the corresponding GT or BLOOM correlation; the GT diagonal is a reference identity. Each BLOOM example uses the observed source and sampled-stage set of its displayed GT comparison. Networks retain the source and its 20 prior targets at fixed positions across GT and BLOOM. Node colours match the scatter key. Coloured arrows encode finite correlations on [−1, 1]; grey arrows and targets retain prior context where no finite association is available. Arrow direction follows the supplied prior. **F,** Expression and prior context for selected genes. The three screens show 25 selected genes per dataset. Columns summarize observed temporal variance, state specificity (standardized across-state mean range), prior outdegree, prior indegree and membership counts in gene programmes curated from the original dataset publications. Colours are average-tie percentile ranks, computed separately for each component over the full 70-, 100- or 135-gene panel before display selection, with limits [0, 1]. Tied ranks can give a zero raw value a nonzero percentile. **G,** Temporal organization of developmental gene-module expression. Target-expression summaries compare GT with BLOOM across the three datasets. Represented target counts are *Runx2*, 15; *Sp9*, 24; *Sp8*, 28; *Sox8*, 22; *Hand2*, 37; *Sall1*, 30; *Prox1*, 41; and *Foxf1*, 72. Scores are weighted means of observed-reference-standardized target expression, using positive absolute prior weights renormalized over represented targets. GT and BLOOM share each dataset’s full-table percentile-based display limits.

Adjacent-stage rank stability captures both sustained gene-expression ordering and intervals of reorganization (Fig. 5a). BLOOM closely follows the major changes in the prenatal and cortical curves, including the pronounced intermediate reduction in rank stability in the second cortical window. Mean absolute differences between the observed and reconstructed rank-stability curves are 0.027 and 0.066 in the two prenatal windows, and 0.036 and 0.061 in cortex. The later endothelial window is also closely followed, with a mean absolute difference of 0.052. Preserving both stable expression ordering and intervals of rank reorganization helps capture the continuity and transitions of developmental gene programmes.

Gene-level timing complements this population-wide summary (Fig. 5b). The paired peak-time and half-range-crossing views retain many of the relative timing relationships among the selected genes. Among the 25 displayed genes, peak times agree exactly for 20 and 19 genes in the two cortical windows, and for 19 and 18 genes in the endothelial windows. Prenatal agreement is more variable, with 7 and 18 exact matches. Half-range crossings provide a second descriptor of expression progression, including cases in which peak positions alone are similar. Together, the rank and descriptor panels support preservation of substantial temporal structure while exposing the windows in which timing is less faithfully reconstructed.

The coexpression matrices extend this analysis from individual trajectories to coordinated expression (Fig. 5c). BLOOM retains recognizable off-diagonal blocks and contrasts among gene pairs in each dataset, alongside changes in association strength. Correlations are calculated after centring expression within each saved age/stage group, reducing the contribution of a shared temporal trend to the displayed associations. This makes the panel complementary to the timing analysis: genes can preserve their temporal maxima while differing in their within-stage coordination. The matrices therefore reveal coordinated gene-expression relationships within developmental stages, providing a molecular context for the reconstructed cell states and their associated programmes.

Paired temporal mean-expression changes provide a direct check on the direction and magnitude of the reconstructed dynamics (Fig. 5d). Across the 25 selected genes and all adjacent saved-stage contrasts, observed and reconstructed changes have Pearson correlations of 0.653 in the prenatal dataset, 0.981 in cortex and 0.902 in STED-EC. BLOOM retains prominent increases and decreases, including the cortical *Car10* increase and the endothelial *Mdk* decline. Prenatal amplitudes are more variable: the *Col2a1* increase between saved stages 23 and 25 and the accompanying *Col1a2* decrease are reproduced with attenuated magnitudes.

We then placed these expression-level results in the context of the supplied regulatory prior (Fig. 5e). The scatterplots compare within-stage residual source–target associations between observed expression and BLOOM, while the circular graphs place those associations within the same prior network.

In the prenatal dataset, *Runx2*, *Pparg* and *Ltf* connect the reconstructed associations to distinct developmental contexts. *Runx2* has strong experimental support in osteoblast differentiation: its disruption prevents normal bone formation [79], and conditional deletion in committed osteoblasts reduces expression of matrix genes including *Col1a1* and *Col1a2* [80]. Both collagen genes occur among the three measured targets in the displayed *Runx2* example, giving the expression comparison a specific skeletal-development context. BLOOM retains their positive association with *Runx2*, with stronger reconstructed correlations than in GT. BLOOM retains the positive *Pparg* – *Car3* association, connecting two experimentally linked components of adipogenic differentiation: depletion of carbonic anhydrase III increases PPAR*γ*2 expression and adipocyte differentiation [81]. The original prenatal atlas nominates *Ltf* as a candidate associated with the emergence of monocytic myeloid-derived suppressor cells [8]; independent lactoferrin perturbation experiments provide a functional neonatal myeloid context [82]. Its displayed associations with *Lyz2*, *Ngp* and *S100a9* are consequently interpretable as a myeloid-associated module.

The cortical examples connect neuronal and oligodendroglial developmental programmes. *Sp8* and *Sp9* have experimentally established roles in CGE-derived cortical interneuron development [68], and *Sp9* also contributes to MGE-derived cortical interneuron development [83]. The displayed *Sp8* –*Htr3a* association therefore has a clear interneuron-associated context. The oligodendroglial examples are particularly informative: the *Olig1* panel includes *Gpr17* and *Pdgfra*, and its two measured associations have a mean absolute GT–BLOOM difference of 0.066. Experiments identify GPR17 as an OLIG1-regulated timing factor: sustained expression inhibits oligodendrocyte maturation, whereas loss of GPR17 accelerates myelination [84].

These associations link the reconstructed cortical profiles to molecular programmes of interneuron development and oligodendrocyte maturation.

The *Tcf7l2* example adds a complementary oligodendroglial differentiation context. Conditional *Tcf7l2* ablation impairs oligodendrocyte differentiation, and the published mutant expression table reports reduced *9630013A20Rik*, one of the targets represented here [85]. This transcript is also known as *lncOL1*, for which genetic and functional experiments establish a contribution to myelination [86]. The other displayed target, *Tns3*, has independent functional evidence in nascent oligodendrocyte differentiation [87]. Their joint inclusion makes the reconstructed expression associations biologically coherent.

The endothelial examples span lymphatic identity, embryonic vascular development and vascular integrity. *Prox1* is required for acquisition of the lymphatic endothelial phenotype [88], providing a developmental context for the displayed *Prox1* –*Lyve1* association. Endothelial-specific *Foxf1* perturbation demonstrates a requirement for embryonic vascular formation and VEGF signalling [73]. The *Foxf1* example includes seven measured targets and has a mean absolute association difference of 0.126, supporting substantial retention of this biologically relevant expression organization. *Hand2* has experimentally established functions within the embryonic endocardium [89]; its selected target associations show larger discrepancies, emphasizing that an established developmental regulator can remain a challenging reconstruction example. The *Klf2* panel includes *Klf4*, linking it to genetic evidence that KLF2 and KLF4 cooperate to maintain embryonic vascular integrity [90]. BLOOM retains the *Klf2* –*Klf4* association, connecting its reconstructed vascular expression profiles to a genetically supported programme of embryonic endothelial integrity.

The expression and prior screens reveal complementary signatures of structural genes and developmental regulators among the selected genes (Fig. 5f). In the prenatal atlas, the collagen genes *Col2a1* and *Col1a2* show pronounced temporal variance, with *Col1a2* also showing prominent incoming prior connectivity. In contrast, *Runx2* stands out for outgoing prior connectivity and programme membership, placing matrix-gene expression alongside a regulator required for osteoblast differentiation and bone-matrix gene expression [79, 80]. In visual cortex, *Sp8* and *Sp9* combine comparatively low temporal variance with strong state specificity and outgoing prior connectivity. Their established roles in interneuron development [68] illustrate how these features retain biological information beyond the magnitude of temporal expression change. In STED-EC, *Prox1* similarly combines low temporal variance with strong state specificity and outgoing connectivity, whereas *Foxf1* shows more prominent temporal variation alongside high state specificity and outgoing connectivity. These profiles provide distinct contexts for lymphatic specification and embryonic vascular formation [73, 88]. Read alongside BLOOM’s reconstructed collagen changes, source–target associations and module trajectories (Fig. 5d,e,g), these complementary signatures connect the recovered expression patterns to both temporally varying effector genes and state-associated developmental regulators.

Finally, covered-target module scores test whether broader expression summaries retain coherent temporal behaviour (Fig. 5g). These scores aggregate 15–72 native-assay target genes per displayed module, extending beyond the few targets in each network inset. Across the four cortical saved stages, the *Sp9* -associated score declines, the *Sox8* -associated score has an intermediate maximum, and the *Sp8* -associated score fluctuates across the saved stages. *Sox8* has an experimentally established contribution to oligodendrocyte differentiation [91], complementing the neuronal context of *Sp8* and *Sp9*. BLOOM retains these distinct temporal shapes, with module-wise Pearson correlations of 0.984 for *Sp9*, 0.997 for *Sox8* and 0.907 for *Sp8*, despite offsets in score amplitude. Across the five STED-EC saved stages, rising *Prox1* - and *Foxf1* - associated scores contrast with declining *Hand2* - and *Sall1* -associated scores; all four reconstructed temporal profiles correlate with GT at 0.952–0.997. The prenatal *Runx2* score shows more moderate agreement, particularly at the terminal saved stage. These module-level results support BLOOM’s preservation of differentiated developmental expression patterns, with the strongest temporal agreement in the selected cortical and endothelial modules. Together, the temporal, coexpression and regulatory views connect BLOOM-generated profiles to coordinated gene programmes associated with skeletal, neuronal, glial and vascular development.

### 2.6 BLOOM reconstructs, interpolates and extrapolates developmental population states

We next examined whether BLOOM could extend a developmental atlas from discrete observations to time-conditioned populations under reverse prediction, interpolation, future rollout and historical rollout (Fig. 6). These settings provide complementary evidence: matched reverse predictions can be compared with measured target populations, whereas the interpolated and extrapolated query populations have no independent experimental counterparts in the supplied evaluation. We therefore assessed the latter through their relationship to observed reference states, their time-dependent population organization and their expression summaries. Observed anchors and boundaries are retained explicitly throughout the figure.

**Fig. 6.**
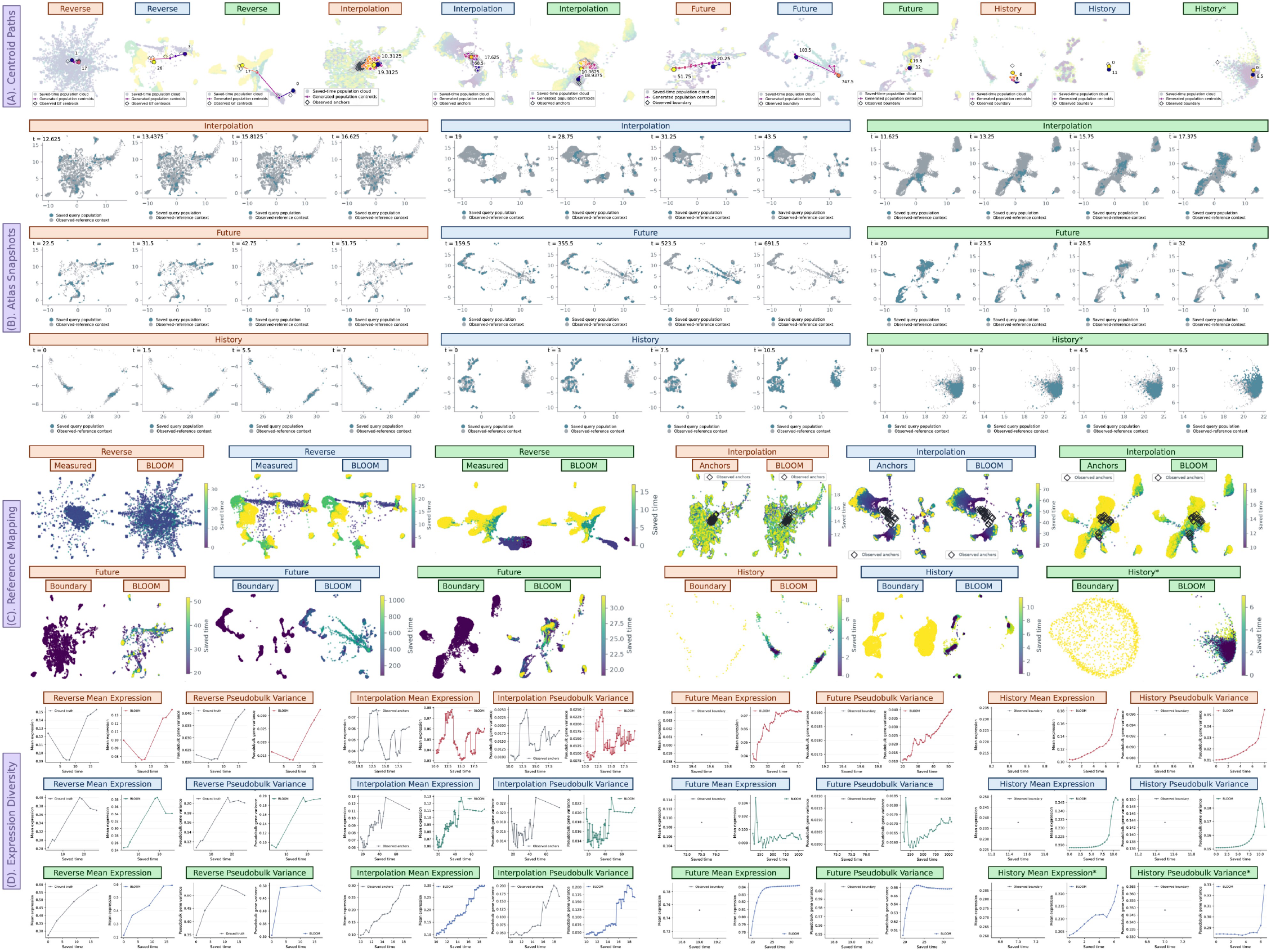
BLOOM reconstructs earlier population states and generates intermediate and extended developmental profiles. Orange, blue and green identify Mouse gastrula-to-birth, Developing mouse visual cortex and STED-EC, respectively. **A,** Population-level transcriptomic paths under reverse reconstruction, interpolation and temporal rollout. Coloured points summarize population centroids in fixed observed-reference UMAP coordinates. Muted clouds provide the corresponding expression context, and diamonds mark observed targets, anchors or boundaries. Paths connect population summaries in saved-time order. **B,** Time-resolved snapshots of interpolated and recursively extended populations. Four selected query coordinates per dataset illustrate interpolation, future rollout and history rollout. Blue points denote the saved query population and grey points show observed-reference context, revealing changes among expression compartments that are averaged within a centroid. **C,** Reference-based placement of reconstructed and simulated populations. Reverse panels compare measured target populations with BLOOM. Interpolation panels show observed anchors and generated intermediate populations. Future and history panels show the measured boundary and the generated continuation. Colours encode the corresponding saved-time coordinate, and interpolation anchors are marked explicitly. **D,** Population-wide expression levels and changes in gene-mean contrast. Global mean expression and pseudobulk gene variance are shown across saved times. The latter is variance across gene-level population means. Reverse axes use target-stage indices; other axes use declared reporting coordinates. Observed and generated subplots retain their exported vertical ranges, so numerical levels are read from their axes. ∗ The asterisk marks the strongly contracted endothelial history example.

Centroid paths, atlas snapshots and reference maps expose different aspects of the generated populations (Fig. 6a–c). Centroids summarize shifts in reference-expression space, whereas the snapshots and maps resolve the separated expression compartments that contribute to those shifts. Across the prenatal, cortical and endothelial settings, generated cells occupy multiple regions of their respective reference atlases. Reading these views together with the population-wide expression summaries allows us to distinguish changes in aggregate expression from redistribution among transcriptomic compartments.

Reverse reconstruction first provides a measured reference for interpreting population-level changes (Fig. 6a,c,d). In the prenatal example, BLOOM retains the reduction in global mean expression around E10.5 and the subsequent increase at E12.75: the observed mean rises from 0.0915 to 0.153, compared with 0.0766 to 0.137 in the reconstructed populations. The cortical reconstruction likewise retains the rise towards P8–P9 and subsequent decrease towards P16, with a P9 mean of 0.386 compared with 0.397 in the measured population. In the cortical reference map, the generated population retains the upper band, the large left-hand compartment and the lower branch, placing these non-monotonic aggregate changes within a structured expression landscape. The endothelial map also retains distinct early- and later-stage regions, alongside the later-stage expression changes. Together, these views show that retrospective inference can recover changes in both expression level and the organization of coexisting developmental states, supporting analysis of earlier population structure from later measurements.

Interpolation resolves changes between measured stages as a sequence of intermediate populations (Fig. 6a–d). The prenatal snapshots remain distributed across a branched reference landscape, while their expression summary rises, falls and rebounds over the interpolated interval. This combination is consistent with a developmental system in which progenitor differentiation, neuronal diversification and erythroid replacement overlap in time, as documented in the original atlas [8]. In visual cortex, the snapshots at reporting coordinates 19, 28.75, 31.25 and 43.5 redistribute query cells among the large upper-left compartment, the lower branch and separated right-hand islands. The cortical mean reaches an intermediate maximum and subsequently settles below that peak. Thus, these interpolations represent changing mixtures of expression states and non-monotonic aggregate profiles rather than a single uniform progression. Such compartment-specific refinement is biologically relevant to the staggered neuronal and glial maturation described across cortical development [9].

The endothelial interpolation provides a particularly detailed example. Its 73 saved population states comprise 19 observed anchors and 54 intermediate queries. Global mean expression changes from approximately 0.0974 to 0.300, while variance across gene-level population means changes from 0.0437 to 0.165, approximately 3.08- and 3.76-fold increases, respectively (Fig. 6d). The variance reaches a local maximum of approximately 0.205 at reporting coordinate 17.875 before declining, while the mean approaches its later plateau. This separates a changing balance among gene means from a simple uniform increase in expression. The snapshots add population context: between coordinates 11.625 and 17.375, query occupancy shifts from the lower diagonal arm towards the broad central-upper compartment while peripheral islands remain represented (Fig. 6b,c). The coexistence and redistribution of these expression compartments are compatible with the combination of shared vascular programmes and organ-associated differentiation documented in endothelial atlases [10, 92]. BLOOM therefore supplies intermediate population states through which the timing of this transcriptomic reorganization can be examined at finer resolution than the measured anchors.

Future rollouts reveal different forms of continuation beyond the measured boundary (Fig. 6a–d). In the prenatal summary series, mean expression initially falls below the boundary, then rises and levels near 0.073, approximately 1.20 times the boundary value of 0.061. Its gene-mean variance continues to rise after the mean has begun to stabilize, indicating increasing contrast among gene-level population means. The prenatal atlas snapshots retain several branches over the illustrated continuation. The endothelial mean instead approaches a plateau near 0.842 from a boundary value of approximately 0.752, an increase of approximately 12%; its gene-mean variance rises initially and then eases from its maximum. Endothelial snapshots retain central and peripheral compartments, with reduced representation of the lower-left arm at later displayed coordinates. In the separate cortical summary series, mean expression remains within a narrow range of approximately 0.097–0.105, compared with a boundary value of 0.109, while gene-mean variance shows an initial decrease followed by partial recovery. This restrained aggregate change is compatible with continuing molecular refinement: the original cortical atlas reports neuronal cluster diversification from 40 at P0 to 93 at P25, with marked changes around eye opening and the onset of the critical period [9]. These distinct rollout shapes provide concrete trajectories for examining population-level remodelling beyond the observation window, without requiring expression to increase uniformly during maturation.

History rollouts extend the analysis before the earliest supplied observation (Fig. 6a–d). In the prenatal summary series, moving backwards from the boundary is accompanied by lower global expression: generated means span approximately 0.10–0.18 compared with a boundary mean of 0.223. The displayed prenatal populations nevertheless retain two narrow expression arms. Cortical history populations also retain separated compartments, while their mean remains within approximately 0.92–1.04 times its boundary value. The preservation of multiple retrospective compartments is biologically meaningful because earlier cortical development contains coexisting progenitor and differentiating populations, with neurogenic and gliogenic programmes unfolding on different schedules [93]. The starred endothelial history run behaves differently: generated cells contract into a compact cloud even though variance across gene means remains substantial. It therefore provides a direct example in which an aggregate expression summary and population organization diverge. The prenatal and cortical cases show how retrospective rollouts can retain a structured mixture of expression states while proposing earlier population configurations from a limited measured window.

Across the four temporal tasks, BLOOM recovers non-monotonic trends in measured reverse comparisons, resolves intermediate changes among coexisting expression compartments, and produces distinct history and future continuations of population-level expression. The contrast between nearly maintained means, changing gene-mean variance and redistributed atlas occupancy shows why developmental extension requires more than a single aggregate trajectory. These results turn sparse atlases into temporally queryable population models, providing intermediate and extended states for analysing developmental reorganization. The following gene-level analyses examine which molecular programmes accompany these population changes.

### 2.7 BLOOM recovers marker trajectories and proposes gene timing beyond observed stages

We next examined whether the population continuations retained interpretable molecular structure at the level of selected developmental genes. Fixed 25-gene expression profiles, individual marker trajectories, adjacent gene-rank correlations and temporal descriptors provided complementary views of this question (Fig. 7). Measured reverse predictions were compared with their matching observed targets. Interpolation instead supplied generated profiles between observed anchors, whereas future predictions extended beyond an observed boundary. We therefore used reverse comparisons to assess measured molecular agreement and used the unmeasured profiles to formulate conditional hypotheses about gene-specific dynamics.

**Fig. 7.**
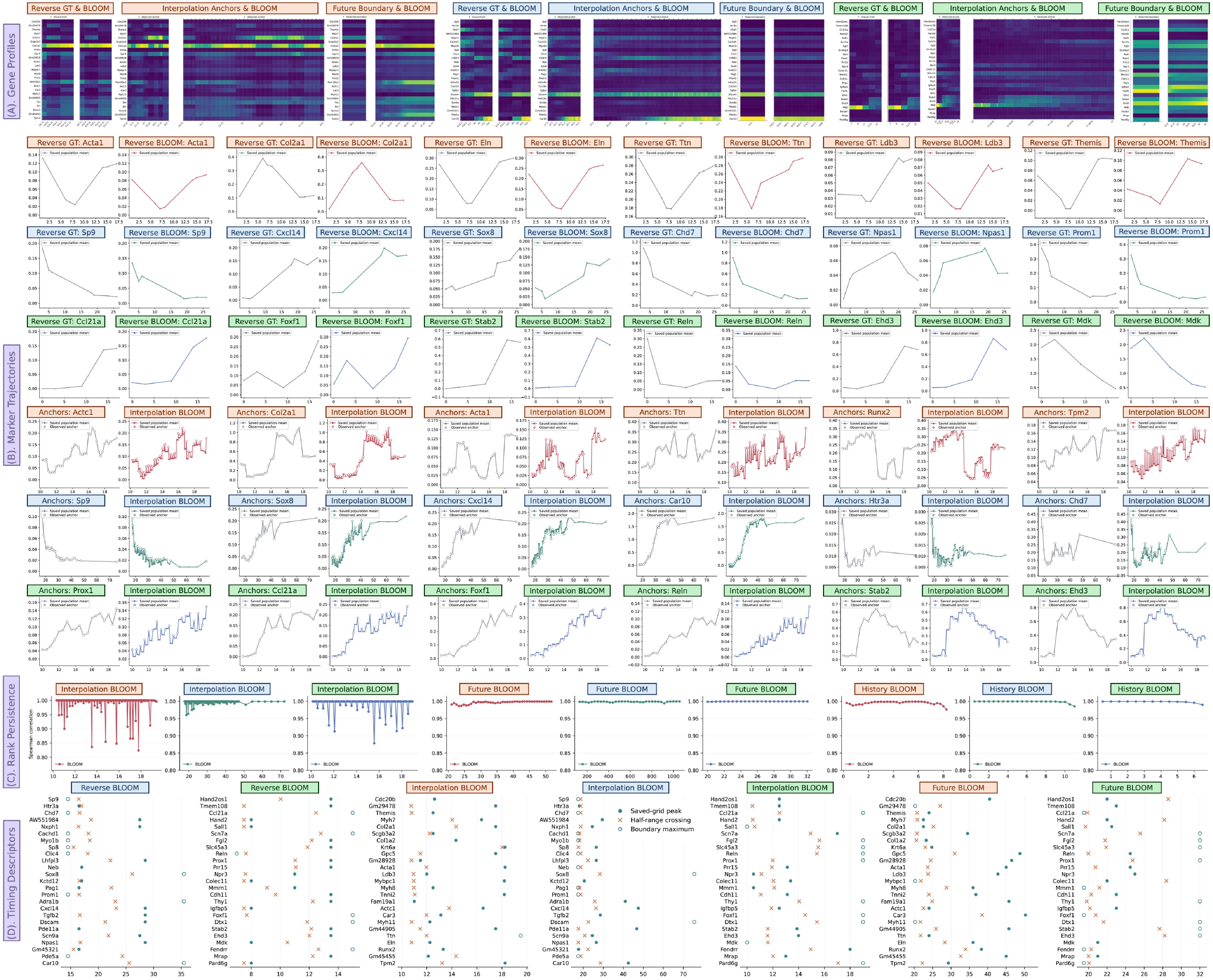
BLOOM reconstructs marker expression and estimates gene-specific temporal landmarks. Orange, blue and green identify Mouse gastrula-to-birth, Developing mouse visual cortex and STED-EC, respectively. **A,** Fixed 25-gene population-mean profiles for measured reverse prediction, interpolation and future rollout. Reference and BLOOM within each pair share a colour scale. **B,** Selected marker trajectories. Grey denotes matching reverse GT or observed interpolation anchors, and coloured traces denote BLOOM predictions. Each subplot retains its displayed vertical scale. **C,** Spearman correlation of gene-wise population means at adjacent saved query times, using cohort-specific exported gene sets. Interpolation excludes anchor rows; the excursions below 0.99 cross omitted-anchor junctions. STED-EC history is diagnostic because high rank persistence coexists with strong cell-variance contraction. **D,** Saved-grid maxima and first half-range crossings for the selected genes. Filled circles denote interior maxima, crosses denote defined crossings and open circles denote boundary maxima. Missing crosses remain undefined; boundary maxima are window-censored.

In the prenatal mouse dataset, the selected traces sampled several biologically distinct developmental compartments (Fig. 7a,b). *Acta1* and *Actc1* are expressed during embryonic striated-muscle development, including coexpression in developing cardiac and skeletal muscle, while *Ldb3* /*Cypher* and *Ttn* relate to sarcomeric structure and maturation [94–96]. The matrix-related genes broadened this assessment: type-II collagen encoded by *Col2a1* is required for normal endochondral skeletogenesis, and elastin contributes to arterial morphogenesis [97, 98]. Across the six displayed reverse markers, descriptive temporal Pearson correlations with the corresponding measured gene means ranged from 0.838 to 0.998 over seven saved target stages. *Acta1*, *Col2a1* and *Eln* achieved correlations of 0.980, 0.943 and 0.998, respectively, supporting recovery of distinct troughs, rises and nonmonotonic profiles rather than a common rescaling of all markers.

The visual-cortex panels similarly preserved contrasting molecular courses (Fig. 7a,b). In measured reverse prediction, *Sp9* and *Chd7* declined across the displayed targets, whereas *Cxcl14* and *Sox8* increased overall; their temporal correlations were 0.988, 0.990, 0.992 and 0.957, respectively. These genes interrogate complementary developmental processes. SP8/SP9 regulate caudal ganglionic eminence-derived cortical interneuron development, and the associated marker context includes *Htr3a* and superficial-layer *Cxcl14* expression [99]. HTR3A also has experimentally established, stage-dependent roles during cortical interneuron invasion, making its changing expression informative without treating it as a monotonic maturity clock [100]. *Sox8* and *Chd7* add regulators associated with oligodendroglial differentiation, while *Car10*, encoding the neurexin-associated protein CA10, provides a separate synaptic adhesion-related molecular context [101–103]. The interpolation examples retained broad decreases in *Sp9* /*Chd7* alongside increases in *Sox8* /*Cxcl14* /*Car10*. The opposing *Sox8* and *Chd7* courses show that genes associated with the same broad developmental compartment can follow different population-mean trajectories, making their individual timing informative alongside a shared oligodendroglial annotation. They do not assign all markers to one interneuron lineage or demonstrate functional synaptic or myelin maturation.

The endothelial dataset provided particularly interpretable contrasts among organ-associated programmes (Fig. 7a,b). *Prox1* and *Ccl21a* provide lymphatic-associated context, and endothelial Reelin has a demonstrated role in collecting lymphatic-vessel development [92, 104, 105]. *Foxf1* is required for embryonic endothelial vascular development, while *Stab2* marks fetal liver-sinusoidal specialization and *Ehd3* is associated with glomerular endothelial differentiation [106–108]. BLOOM reproduced measured reverse courses for these distinct readouts: the displayed *Ccl21a*, *Stab2* and *Ehd3* traces had temporal correlations of 0.992, 0.997 and 0.993 across five target stages. The interpolation profiles then separated broadly increasing *Prox1* /*Ccl21a*/*Foxf1* /*Reln* from the intermediate-window maxima and subsequent decreases of *Stab2* /*Ehd3*. This pattern is consistent with differently timed organ-associated expression programmes within an endothelial population whose tissue-dependent heterogeneity is independently established [92]. It motivates subtype-resolved analysis rather than a liver-to-kidney-to-lymphatic lineage sequence.

Adjacent gene-rank correlations supplied a complementary continuity diagnostic (Fig. 7c). In the displayed future rollouts, median Spearman correlations between neighbouring population-mean gene vectors were 0.9985, 0.9993 and 0.9999 for the prenatal, cortical and endothelial datasets, respectively. Thus, substantial gene-specific mean-expression changes could occur while the broader ordering of expressed genes remained largely persistent. The most conspicuous interpolation excursions required a more specific interpretation: every interval with correlation below 0.99 crossed a junction between query segments separated by an omitted observed anchor. These dips locate sensitivity at anchor junctions; they do not independently establish biological regulatory switches. Likewise, the STED-EC history example retained high rank correlation despite severe cell-level variability contraction and is explicitly marked as diagnostic. Rank persistence therefore describes one aspect of the generated molecular organization and complements, rather than replaces, expression-amplitude and dispersion checks.

The temporal descriptors made selected aspects of this molecular structure directly inspectable (Fig. 7d). A saved-grid maximum summarizes the largest population mean within the available window. The first half-range crossing uses each gene’s own minimum and maximum and follows the direction of its endpoint change, so it can describe either an increase or a decrease. In measured reverse prediction, the maximum stage matched GT for 19 of the 25 selected genes in both cortex and STED-EC. In cortex dataset, *Sp9*, *Chd7* and *Prom1* had their largest saved values at E14.5, whereas *Sox8* and *Car10* had their largest values at P16; these maximum stages matched their observed references. In STED-EC, the matched *Stab2* /*Ehd3* maxima occurred at E13.5, whereas *Ccl21a*/*Foxf1* maxima occurred at E15. This constitutes measured support for specific marker-dependent timing differences.

For interpolation, the descriptor panel identified candidate landmarks within the conditioned expression paths. Prenatal *Runx2*, an essential osteoblast-differentiation regulator [109], had an interior query-grid maximum near reporting coordinate 13.31, distinct from the observed-anchor maximum of *Actc1* at 16.50 and the terminal-window maximum of *Ttn* at 19.50. In STED, the *Stab2* maximum near 13.88 occurred at an intermediate query, whereas the *Ehd3* maximum at 14.00 coincided with an anchor; their half-range crossings were both near 11.6, compared with approximately 15.4 for *Reln*. These differences provide concrete candidates for testing the relative timing of distinct tissue-associated programmes. Their evidential status remains conditional: 18/25 prenatal, 22/25 cortical and 21/25 endothelial interpolation maxima occurred at supplied anchor times. The figure therefore distinguishes additional model-derived query landmarks from timing inherited through the observed context. Neither the row order nor the crossing-to-maximum relationship defines a causal regulatory cascade.

The future descriptors extended this analysis into experimentally unmeasured contexts. In the prenatal dataset, *Ttn* and *Runx2* showed distinct interior maxima at reporting coordinates 24.00 and 38.25, respectively, proposing different continuation kinetics for sarcomeric and osteogenic readouts. In STED-EC, *Foxf1* decreased from 0.324 to 0.241 across the saved future queries, while *Stab2* increased from 0.225 to 0.297 and *Ehd3* from 0.347 to 0.399. Given the different vascular-specialization contexts of these markers [73, 107, 108], this opposing behaviour is a testable hypothesis of selective changes in specialization-associated markers. In postnatal mouse kidney, EHD3 protein is detected in capillary-loop and mature glomeruli but not earlier comma- and S-shaped structures [108], providing a maturation-associated context for testing the predicted *Ehd3* increase. *Prox1*, *Stab2* and *Ehd3* reached their largest saved values at the final STED-EC coordinate, so their maxima remain window-censored. The much smaller ranges of *Ccl21a* and *Reln* provide persistence context and do not justify precise future peak dates. Population composition and within-state regulation could both contribute to these aggregate predictions, and some low-expression queries retain negative saved means. With these limits explicit, BLOOM supplies experimentally addressable, gene-specific hypotheses beyond measured stages alongside its demonstrated recovery of selected reverse marker trajectories.

### 2.8 BLOOM reveals biologically interpretable developmental responses to single-gene input edits

To investigate whether BLOOM’s developmental representations encode interpretable gene dependencies, we examined predictions following single-gene edits to the saved source history across the prenatal, cortical and endothelial atlases (Fig. 8). In each experiment, one gene coordinate was set to zero in the final source-history frame, while the fitted model and target conditioning were retained. The matched baseline was the same model’s unedited prediction. Across the 12 edited-gene examples in panel a, all 78 saved-stage comparisons showed lower predicted values of the edited coordinate after the input edit. This consistent self-response establishes that the edited context influences subsequent predictions and motivates examination of the accompanying secondary responses.

**Fig. 8.**
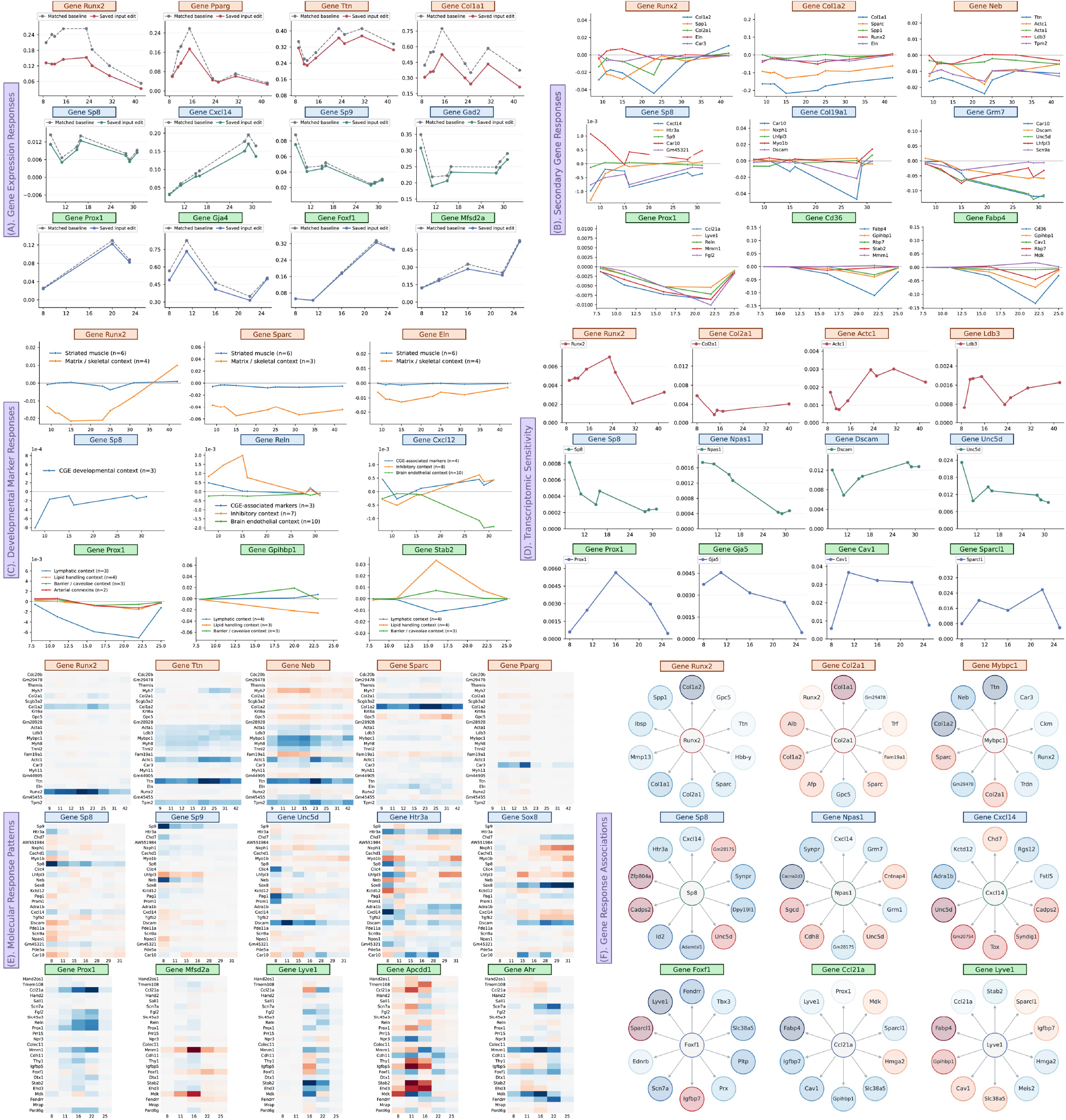
BLOOM predicts context-dependent responses to single-gene source-history edits. Orange, blue and green identify Mouse gastrula-to-birth, Developing mouse visual cortex and STED-EC, respectively. Horizontal coordinates are saved target-stage indices. **A,** BLOOM propagates single-gene input edits into subsequent expression predictions. One gene coordinate is set to zero in the final source-history frame, with the model and target conditioning retained; generated outputs are not clamped. Curves compare populationmean expression of the edited gene between matched unedited and edited inputs, showing consistently reduced predictions after the input change. **B,** Single-gene edits elicit selective responses in associated genes. Lines show edited-minus-baseline population-mean differences for selected other genes across target stages. Examples include reduced *Spp1* after *Runx2* editing, reduced *Cxcl14* after *Sp8* editing and reduced lymphatic-marker expression after *Prox1* editing. **C,** Marker-set responses connect input edits to developmental programmes. Curves summarize mean expression differences within literature-informed marker sets, excluding the edited gene and requiring at least two covered members; *n* gives the number of included markers. The *Prox1* and *Gpihbp1* examples show coherent decreases in lymphatic- and lipid-associated expression contexts, respectively. **D,** Transcriptomic response magnitude varies with the edited gene and target stage. Curves show root-mean-square expression differences across the remaining exported genes after excluding the edited coordinate. These profiles resolve stage-dependent sensitivity, including decreasing cortical *Sp8* responses and non-monotonic *Dscam* responses, on each component’s native expression scale. **E,** Heatmaps resolve signed molecular responses across developmental stages. Rows show fixed 25-gene response panels and columns show saved target stages as categorical coordinates; colours encode edited-minus-baseline population means. Red and blue denote positive and negative responses, respectively, on each panel’s own display scale, revealing selective changes among associated genes. **F,** Association diagrams summarize the molecular context of selected input edits. Spokes connect the edited input to responding genes, with node colour showing each gene’s signed response at its own stage of maximum absolute change (red, positive; blue, negative). These model-derived associations include *Col2a1* –*Col1a1*, *Sp8* –*Cxcl14* and *Foxf1* –*Fendrr*.

In the prenatal atlas, several responses were directionally compatible with established skeletal developmental dependencies. Following the *Runx2* input edit, *Spp1* decreased at all eight saved target stages and *Col1a2* decreased at seven (Fig. 8b). These directions are consistent with the loss of osteopontin expression in immature osteoblasts of *Cbfa1* /*Runx2* -null embryos and reduced matrix-gene expression after inhibition of CBFA1 activity in differentiated osteoblasts [110, 111]. The matrix-associated marker-set mean was also negative at seven stages, with a terminal positive response indicating temporal selectivity rather than uniform suppression of all matrix components (Fig. 8c). A complementary association emerged after the *Col2a1* edit: *Col1a1* increased at all eight stages, with its largest saved difference at stage 31 (Fig. 8f). Embryonic *Col2a1* -null cartilage ectopically expresses type-I collagen, providing a specific experimental precedent for this response direction [112]. This correspondence motivates a matrix-substitution hypothesis and illustrates how BLOOM can generate testable predictions of extracellular-matrix remodelling.

The muscle-associated probes revealed coordinated, but selective, responses among structural genes. Editing *Neb* reduced *Ttn*, *Actc1*, *Acta1* and *Tpm2* at all eight saved targets (Fig. 8b). Nebulin deficiency experimentally alters thin-filament structure and contractile function, supporting the functional relevance of this sarcomeric context without independently establishing the predicted transcript directions [113]. More specifically, *Ldb3* editing reduced *Acta1* and *Actc1* at all eight stages, concordant in direction with reduced expression of these actin transcripts in neonatal mouse hearts lacking long Cypher isoforms [114]. These selected agreements show that BLOOM can express biologically relevant responses beyond attenuation of the edited coordinate itself.

The cortical atlas provided a specific literature-grounded developmental comparison. Following *Sp8* editing, *Cxcl14* decreased at every one of the seven saved target stages (Fig. 8b,f). This agrees with experimental evidence that cortical *Cxcl14* expression depends on SP8 in the context of caudal ganglionic eminence-derived interneurons [115]. The self-excluded mean of the displayed *Htr3a*/*Sp9* /*Cxcl14* marker set was negative at all seven stages (Fig. 8c), extending the single-gene observation to a developmentally relevant contextual summary. Individual members nevertheless differed: *Htr3a* decreased at the first four stages and showed small positive changes later, while *Sp9* changed sign. The result therefore supports selective sensitivity within an interneuron context rather than complete suppression of a lineage programme. The additional *Col19a1* and *Grm7* probes address genes with established roles in inhibitory synapse assembly and the frequency-dependent recruitment of somatostatin interneurons, respectively [116, 117]. Their secondary transcript responses remain predictions for experimental testing.

The endothelial atlas linked input sensitivity to lymphatic identity and vascular metabolic context. The *Prox1* edit in the selected secondary-response example reduced *Ccl21a* and *Lyve1* at all five saved stages, with the largest decreases in both genes at stage 22 (Fig. 8b). These directions agree with genetic evidence that *Prox1* is required for the acquisition and maintenance of lymphatic endothelial identity, including expression of LYVE1 and the CCL21/SLC marker [118, 119]. The corresponding self-excluded lymphatic-context mean, comprising *Ccl21a*, *Lyve1* and *Reln*, was negative at all five stages; all 15 member-by-stage responses shared this sign (Fig. 8c). Reelin has also been experimentally implicated in lymphatic vessel formation, supporting its inclusion in this contextual marker set [120]. Unlike the mixed *Htr3a*/*Sp9* /*Cxcl14* responses to *Sp8* editing, the uniformly negative lymphatic-marker responses identify a coordinated predicted shift across the selected identity-associated genes.

The lipid-associated responses identified another interpretable endothelial hypothesis. Editing *Cd36* reduced *Fabp4* at all five stages; editing *Fabp4* reciprocally reduced *Cd36* and also lowered *Gpihbp1* (Fig. 8b). In the marker-set analysis, the *Gpihbp1* edit reduced the *Cd36* /*Fabp4* /*Rbp7* mean at all three exported stages, with concordant signs in all nine member-by-stage responses (Fig. 8c). Endothelial PPAR*γ*-dependent coordination of CD36 and FABP4, the contribution of endothelial CD36 to tissue fatty-acid uptake, and GPIHBP1-mediated transport of lipoprotein lipase establish a common lipid-handling context for these genes [121–123]. RBP7 additionally links the marker set to PPAR*γ*-dependent endothelial antioxidant protection through adiponectin [124]. The association diagrams additionally identified a decrease in the long noncoding RNA *Fendrr* after *Foxf1* editing (Fig. 8f), matching the direction reported following endothelial *Foxf1* deletion [125]. Other arrows, including responses to *Ccl21a* or *Lyve1* edits, remain prospective associations. In particular, normal lymphatic development in *Lyve1* -deficient mice argues against interpreting the displayed *Lyve1* responses as proof of an indispensable upstream lymphatic-specification function [126].

Broader expression summaries resolved the timing and molecular breadth of the predicted responses. Downstream root-mean-square responses, calculated after excluding the edited gene, varied across saved stages (Fig. 8d). For example, the cortical *Sp8*, *Npas1* and *Unc5d* examples had lower sensitivity at their final than initial stage, whereas *Dscam* was nonmonotonic. These curves describe model sensitivity within each exported gene universe, rather than gene essentiality or comparative perturbation accuracy. The 25-gene heatmaps preserve signed response heterogeneity, while the association diagrams summarize each gene at its own strongest-response stage (Fig. 8e,f).

Together, these analyses show that BLOOM links single-gene input changes to selective, stage-dependent responses across skeletal, muscular, neuronal and vascular expression programmes. Concordant examples, including *Runx2* –*Spp1*, *Sp8* –*Cxcl14*, *Prox1* -associated lymphatic markers and *Foxf1* –*Fendrr*, place the predicted responses in established developmental contexts. These in silico experiments extend the transcriptomic virtual embryo from temporal simulation to hypothesis generation, helping prioritize gene dependencies, molecular programmes and developmental stages for targeted experimental investigation.

## 3 Discussion

BLOOM provides a transcriptomic framework for studying development as a sequence of heterogeneous cell populations that can be queried across time. By combining an observed population history with inferred cell correspondences and developmental, anatomical and regulatory knowledge, it connects temporal prediction with the biological organization represented in developmental atlases. Across three mouse datasets, the leading gene- and cell-expression correlations in comparisons with 22 time-series forecasting baselines were accompanied by recognizable population structure, marker distributions and programme-associated expression. These findings support the value of evaluating a developmental generator at several resolutions: agreement in expression patterns becomes more informative when the generated populations also retain distinctions among cell states and coordinated molecular programmes.

A central feature of BLOOM is the integration of temporal history and biological knowledge within the representation that conditions generation. Recent approaches have advanced developmental modeling through molecular kinetics, temporally coherent flow learning and sequence-based diffusion generation [22, 24, 28]. BLOOM brings an ordered window of populations together with relationships among cells, developmental states, tissues and genes, using this context to condition both a biologically informed Gaussian base and the flow-matching generator. This design makes biological resources available throughout conditional expression generation.

The biological analyses illustrate why developmental prediction benefits from information beyond the magnitude of temporal expression change. In the expression and prior screens, cortical *Sp8* /*Sp9* and endothelial *Prox1* combined modest population-wide temporal variation with strong cell-state specificity and outgoing prior connectivity. Their relevance therefore lies partly in the populations and molecular relationships with which they are associated. Reconstruction of interneuron-associated markers, oligodendroglial modules and organ-associated endothelial signatures shows how generated expression can remain interpretable at these different biological scales. Coexpression and source–target association analyses add information about coordinated expression within stages, while programme trajectories describe changes in the expression of their constituent genes over time. Together, these views connect predictive agreement to specific developmental contexts and identify molecular readouts for subsequent investigation.

The temporal applications give this representation a practical role between atlas construction and experimental design. Earlier-stage reconstruction provides a measured test of retrospective inference, whereas interpolation and recursive history and future rollouts propose populations between and outside the supplied observations. The different predicted courses of prenatal *Ttn* and *Runx2*, and the contrasting endothelial *Foxf1*, *Stab2* and *Ehd3* continuations, illustrate how temporal extension can generate hypotheses about differently timed molecular programmes. Such predictions can help select developmental intervals and markers for additional sampling. Tissue- and cell-state-resolved measurements would then distinguish expression changes within a population from changes in the mixture of populations contributing to an aggregate trajectory.

Reliable temporal extension will require explicit assessment of predictive diversity and uncertainty. In the endothelial history example, high persistence of gene rankings coexisted with substantial contraction of cell-level variability. Smooth population means or stable expression ranks can therefore conceal loss of cellular heterogeneity as generated populations increasingly replace measured inputs. Future evaluation should measure dispersion, state occupancy and distributional agreement as a function of rollout horizon, alongside independent temporal tests to calibrate uncertainty. Interpolated landmarks likewise require denser measurements: many selected expression maxima occurred at supplied anchors, and terminal maxima in future rollouts remain window-censored. Establishing reliable horizons and uncertainty for both population structure and gene timing would make these extensions more useful for planning experiments.

Single-gene input edits extend BLOOM from temporal queries to the prioritization of developmental response hypotheses. The *Runx2* –*Spp1*, *Sp8* –*Cxcl14*, *Prox1* - associated lymphatic-marker and *Foxf1* –*Fendrr* responses connect model sensitivity to established developmental contexts. However, the current perturbation workflow is limited to single-gene edits in the final source-history frame and lacks matched experimental perturbation outcomes. These edits do not impose persistent depletion throughout the generated course. Specific disagreements with published experiments define useful targets for refinement: the selected *Actc1* edit decreased *Acta1*, whereas *Actc1* -null newborn hearts showed increased skeletal-actin transcripts [127]; the selected *Ldb3* edit decreased *Myh7*, whereas neonatal mouse hearts lacking long Cypher isoforms showed increased *Myh7*, despite decreases in *Acta1* and *Actc1* [114]. Tissue composition, developmental timing, isoform specificity and intervention persistence are possible contributors to these differences. Paired control and perturbed populations sampled over time would enable direct tests of response direction, magnitude and duration, and support extensions to sustained, graded and combinatorial perturbations.

Multimodal measurements offer a further opportunity to connect temporal expression prediction with additional regulatory and phenotypic information. BLOOM currently models scRNA-seq without directly observing chromatin accessibility or protein abundance. Joint profiling of RNA and chromatin has identified regulatory accessibility changes that precede associated transcriptional changes during differentiation [128], while joint RNA–protein assays provide antibody-defined protein markers alongside transcriptomes [129]. Incorporating these measurements could refine regulatory context and allow evaluation of predicted changes across molecular layers. Spatial measurements could additionally anchor the anatomical context to observed tissue organization.

All three evaluations concern mouse development, so transfer to human systems remains an important next step. Human fetal atlases already provide extensive coverage of cellular diversity across organs [130]; the need is for sufficiently dense temporal sampling, consistent developmental annotations and appropriate perturbation measurements for human prediction and validation. Combining such resources with human organoid time courses could support studies of disease-associated developmental deviations and treatment responses.

By bringing temporal population modeling and biological knowledge into a shared generative framework, BLOOM advances the use of developmental atlases as queryable transcriptomic models. Its measured-stage reconstructions establish a foundation for exploring intermediate states, extended expression trajectories and gene-dependent responses. With broader molecular measurements and prospective temporal and perturbation validation, this framework could help focus experiments on the stages, populations and molecular programmes most informative for understanding development and its disruption.

## 4 Methods

### 4.1 Datasets, expression panels and notation

We analyzed three published mouse developmental resources: the Mouse gastrula-to-birth atlas [8], the Developing mouse visual cortex atlas [9], and the multi-organ endothelial developmental atlas STED-EC [10]. Original developmental-age, cell-state and available anatomical annotations were retained. The highly variable gene (HVG) and author-derived panels were kept separate: the HVG panel contained 200 genes per dataset; the retained author panels contained 70, 100 and 135 genes for prenatal development, visual cortex and STED-EC, respectively. Author candidates were obtained from deduplicated supplied programme–gene memberships and aligned to the available measured genes.

Let *T* denote the number of measured stages, *r* ∈ {1*, . . ., T* } a stage index, and *t_r_* its numerical developmental age. The observed population is *X_r_* ∈ R*^Nr^* ^×^*^G^*, with *N_r_* cells and *G* panel genes. Its *i*th row is *x_r,i_* ∈ R*^G^*, and *X_r,ig_* is gene *g* in that cell. The set of panel genes is, with = *G*. Predictions are denoted *X_r_* and *x_r,i_*. We use *K* for input-history length, *L* for decoder label length, *P* for direct output length, and *H* for the number of recursive rollout steps. Developmental direction is *d* ∈ {−1, +1}; *d* = +1 predicts from earlier stages and *d* = −1 reconstructs from later stages.

### 4.2 Expression preprocessing and temporal transport geometry

Preprocessing used Scanpy [131]. Cells expressing fewer than 200 genes and genes detected in fewer than three cells were removed. Library-size normalization and logarithmic transformation were defined by

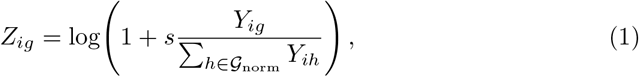

where *Y* is source expression, *s* is the target library size and _norm_ is the gene universe present at normalization.

The prenatal workflow began with a supplied 10,000-gene matrix and additionally retained cells with more than 200 and fewer than 2,000 detected genes and mitochondrial fraction below 5%. Counts were normalized with *s* = 10^4^ and logarithmized before subsetting to 2,000 HVGs selected from preserved counts. Library size and mitochondrial fraction were regressed from a separate layer, which was scaled to unit variance and clipped at 10 before PCA.

Ordinal stage indices, checkpoint time features and reporting coordinates were retained separately. Where embryonic and postnatal labels were combined for reporting, embryonic day *a* mapped to *a* and postnatal day *a* to 19.5 + *a*. Downstream conditioning retained the checkpoint’s time representation and normalization.

### 4.3 Temporal correspondences and pseudo-trajectories

We used moscot [51] to infer transport couplings between adjacent developmental stages from PCA coordinates. Squared Euclidean costs were scaled by their stage-pair mean. Couplings were aligned to original cell identifiers; forward candidates came from coupling rows and backward candidates from the transposed coupling. In the filtered-edge configuration, candidate sets combined the top-*k* links with the smallest descending prefix reaching cumulative mass fraction *q*, and retained weights were renormalized. Paths were extended towards earlier and later stages, stitched in stage order and retained when they covered the required stages. These correspondences supplied both baseline time series and BLOOM trajectory hints.

### 4.4 Regulatory and author-derived biological resources

Expression-derived TF–target associations were inferred with GRNBoost2 in the pySCENIC workflow [52, 53, 132], using preprocessed expression and cells sampled within developmental-stage and cell-type strata. The inference gene set included selected expression genes and available expressed transcription factors. Weighted associations were aligned to the model gene panel for biological conditioning.

Published annotations were converted into directed developmental graphs, cell-to-node assignments and state-associated gene programmes. Primary graph edges had unit weights. For visual cortex, Supplementary Table 5 defined cluster–age relationships, and Supplementary Tables 6 and 9 supplied expression and chromatin-supported gene programmes. Absolute author scores were maximum normalized within module context, with inverse-square-root source-rank weights used when scores were unavailable. State–gene weights combined the two evidence sources equally, *ψ_ug_* = 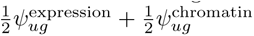, with absent evidence assigned zero. Chromatin-supported memberships entered as fixed published annotations.

For the prenatal atlas, developmental-progression edges defined the primary graph, and original cell assignments linked cells to graph nodes. Supplementary Tables 23 and 24 supplied transition-associated TF and gene lists. Genes in the selected transition direction received inverse-square-root rank weights, additionally scaled by normalized effect-size and adjusted-*P* evidence when per-gene statistics were available. Statistical ranking used increasing adjusted *P*, followed by decreasing absolute effect; TF multipliers were one, with a configurable multiplier for other genes. TF-only and combined programmes were exported separately and attached to target states.

For STED-EC, author-adjusted lineage sequences defined developmental state– period graphs. Cells were mapped to matching annotated nodes, and published state and maturity markers supplied equally weighted gene programmes at compatible nodes.

### 4.5 Time-series baselines and target-stage partitions

Each pseudo-trajectory supplied a multivariate series with genes as variables. If *j_q_*(*r*) identifies its cell at stage *r*, the encoder and decoder inputs were

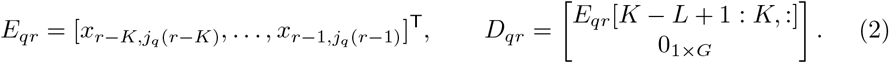

The target was *x_r,jq_* _(_*_r_*_)_. The baseline records used *K* = 8, *L* = 4 and *P* = 1, so an *H*-step forecast requires *H* recursive applications. Models were imported through Time-Series-Library [133]; their final output position supplied the predicted profile.

The baseline set comprised Transformer [48], Informer [35], Reformer [41], Autoformer [29], FEDformer [32], Nonstationary Transformer [38], Crossformer [30], PatchTST [40], iTransformer [50], PAttn [39], TimeXer [46], TimeFilter [45], FiLM [33], TiDE [44], DLinear [31], LightTS [36], TimesNet [47], FreTS [34], TSMixer [43], WPMixer [49], SegRNN [42] and Mamba [37].

Trainable baselines minimized trajectory-occurrence-weighted MSE with Adam, gradient clipping and validation-based checkpoint selection, using float32 and seed 42. The prenatal baseline workflow sampled 10% of trajectory rows; the visual-cortex and STED-EC workflows used their full trajectory tables. Repeated predictions for the same target cell were averaged before unique-target evaluation.

Target stages were assigned to training, validation and test sets. In the random-stage protocol, test stages were sampled first and validation stages were preferentially selected before test stages, using nominal train and validation ratios 0.7 and 0.1 and split seed 42. Only targets with sufficient history were eligible.

### 4.6 Population-history encoding and adaptive biological priors

BLOOM summarizes a directional window *_r,d_* = (*r dK, . . ., r d*), ordered towards the requested target. A shared two-layer GELU cell encoder and masked mean pooling produce stage representations

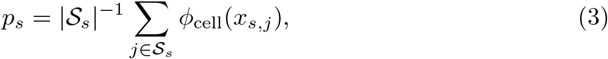

where *_s_* is the seed-fixed source-cell sample. The source proportion of the requested cell state is appended before stage projection, learned window-position embeddings and a single-layer GRU. The soft-conditioning configuration samples cells across the source-stage population. The final temporal state, target-state embedding, encoded trajectory hint *a_r,i_* and its availability indicator form *h*_0_. Hints average linked predecessor occurrences for prediction and successor occurrences for reconstruction. Requested row counts, cell states and available target-indexed annotations are supplied conditioning inputs.

The adaptive model encodes six prior groups: developmental age, cell-identity hierarchy, developmental graph, anatomical/numerical context, gene programmes and GRN. Age features summarize standardized source and target times and their gaps. Categorical features use embeddings, while numerical features retain missingness masks. Graph features combine a learned node embedding with normalized depth, degrees, ancestor/descendant counts and terminal-node status. Programme weights are aligned to model genes and maximum-absolute normalized within states. With regulators as rows and target genes as columns, the retained GRN weights define

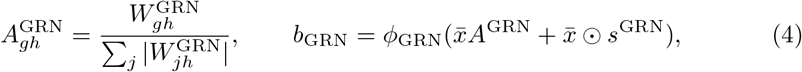

where *x̄* averages source-stage expression means, zero-weight columns remain zero and *s*^GRN^ is the vector of incoming absolute edge weights, maximum normalized across target genes. This support includes retained outside-panel regulators; their expression is not imputed.

For *Q* enabled prior groups, feature encoders produce *b_q_* and learned conditional gates integrate them as

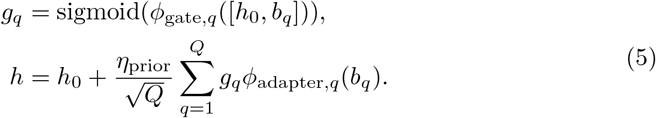

Adapter output layers are zero initialized. The prior schedule controls *η*_prior_ during training, with *η*_prior_ = 1 at inference.

### 4.7 Conditional flow matching, losses and expression readouts

Following flow matching [18], the biologically informed conditional representation defines a diagonal Gaussian base. Its mean and clipped log variance are linear projections of context:

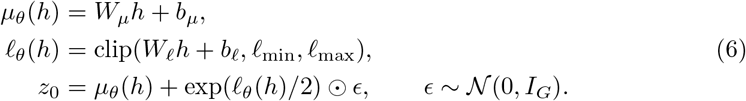

For target endpoint *u_r,i_*= *x_r,i_*, training samples *τ* ∼ Uniform(0, 1) and minimizes

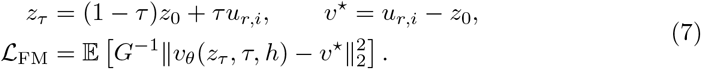

The velocity MLP combines projections of expression, flow coordinate and context. The base remains differentiable through the interpolation path and velocity target. A deterministic head predicts 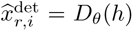, with auxiliary loss 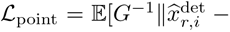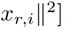. The full adaptive training recipe uses

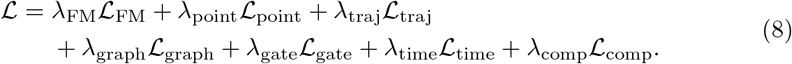

Graph regularization penalizes edge-weighted squared differences between node embeddings. Continuous-time augmentation uses the expression proxy (1 − *f*)*a_r,i_* + *fx* at a sampled source–target fraction *f* ∼ Uniform(0.05, 0.95), reusing the configured expression objective without graph/gate penalties. A separate composition head predicts target-state probabilities from population history and time/GRN context, omitting target-specific state embeddings, trajectory hints and cell-indexed annotations; it uses cross-entropy against training target types. Trajectory hints enter the context without an additional trajectory-attraction penalty in this recipe.

For each of *S* independent base draws, *J* updates integrate the field to give *x*^flow^ = *z_J_* ; the blended prediction is

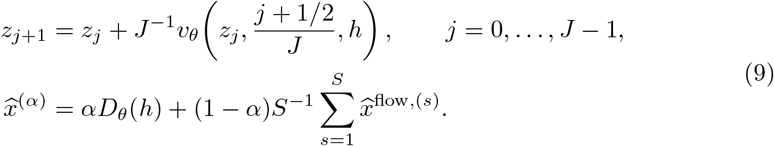

### 4.8 Bidirectional modeling and optimization

Unidirectional BLOOM predicts from earlier populations, and a separately trained reverse control reconstructs from later populations. Bidirectional BLOOM shares the population encoder, prior encoders and generative heads. Direction enters before prior fusion through

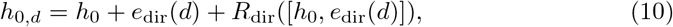

where the direction embedding and optional adapter output are zero initialized. One target-stage partition governs both directions, with direction-specific context eligibility and macro-averaged validation scores.

The supplied training recipes specify hidden width 256 for the population, temporal and flow networks, state-embedding width 32, trajectory-encoder width 128 and prior embedding/hidden widths 16/64, with up to 2,048 source cells per stage. AdamW uses batch size 128, learning rate and weight decay 10^−4^, and gradient clipping at 1. Prior gates start at 0.1, with prior dropout 0.1, five warmup epochs and a ten-epoch ramp. Gaussian log variance is clipped to [−8, 3]; the inference recipe uses eight flow steps and four initial draws. Multi-readout training retains a validation-best checkpoint for each requested readout, with early stopping reset when any readout improves. Test evaluation follows checkpoint selection.

### 4.9 Measured prediction, interpolation, recursive rollouts and single-gene input edits

Downstream analyses reused trained checkpoints without parameter updates. Later-stage prediction and earlier-stage reconstruction used their corresponding temporal direction and retained the checkpoint’s gene order, expression representation and prediction readout.

For interpolation between adjacent measured stages, the requested checkpoint time coordinate was *t_r,f_* = (1 − *f*)*t_r_* + *ft_r_*_+1_, with 0 *< f <* 1. Observed context ended at the left anchor; the right anchor supplied population-size, cell-state and cell-indexed prior templates. Target-time features and time gaps changed across fractions, with the random seed held fixed within each interval.

Recursive future and history rollouts began with the final *K* observed stages in ascending time order and the first *K* observed stages in descending time order, respectively, where *K* is the active context length. Each generated population entered the context window and supplied the next trajectory hint; the window retained its last *K* populations in the requested direction. Target row counts, state labels and cell-indexed annotations were inherited from the corresponding boundary stage throughout the rollout. Time features and expression-derived GRN context updated with the simulated history. Query times followed each component’s reporting grid while retaining checkpoint time normalization.

For single-gene input edits, the selected gene was set to zero in the final source frame and its trajectory hint, in model-input units. Baseline and edited calls shared the checkpoint, target templates, other annotations, readout and random seed. The response of gene *g* to editing gene *j* at stage *r* was

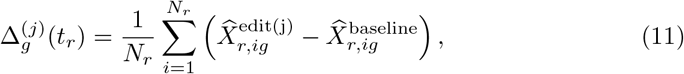

where *N_r_*is the number of evaluated target rows.

### 4.10 Numerical evaluation of measured-stage predictions

Numerical comparisons were performed separately for each dataset using measured forward predictions and their matching observed expression. Let *X_r_, X_r_* ∈ R*^Nr^* ^×^*^G^* denote observed and predicted expression at target stage *r* ∈ R, where *R* = |R|. Their row-stacked matrices are *X, X* R, where *N* = *_r_*_∈R_ *N_r_*; these quantities refer to the evaluation cohort of the model being scored.

#### Cell and gene correlation

For equally sized vectors, Pearson correlation and uncentered cosine similarity were defined as

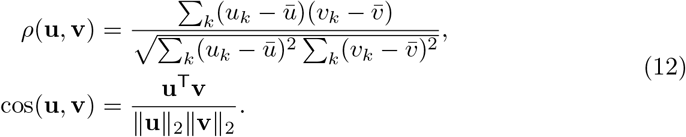

Within each stage, cell correlations compared expression profiles across genes, whereas gene correlations compared expression across matched cells. Denote these by

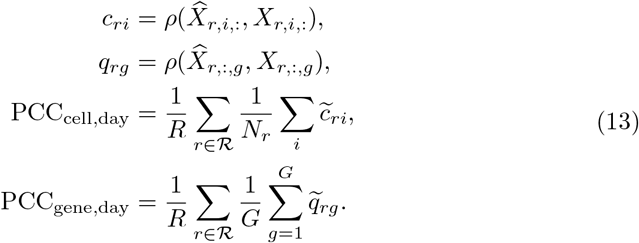

Tildes denote the recorded aggregation convention in which undefined component PCCs contribute zero. Pooled cell and gene PCC means and medians instead summarize row and column correlations of *X* and *X*. Mean row cosine averages defined uncentered similarities, 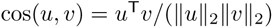, across pooled target rows.

#### Expression agreement and population means

Pooled errors and the global coefficient of determination are

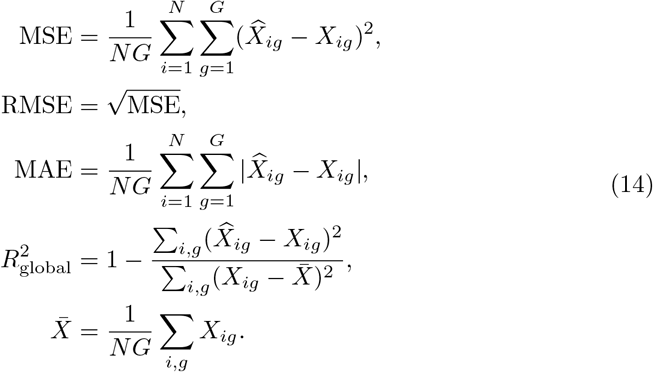

Pseudobulk profiles were arithmetic population means on the saved expression scale. Let 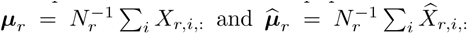, and let ***µ*** and ***µ***^ denote cos(***µ***^, ***µ***); pseudobulk MAE was 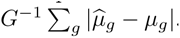. Mean-day pseudobulk PCC averaged defined values of ρ(^ μ*_r_*,μ*_r_*) equally across target stages, with valid-stage coverage recorded. Time–gene pseudobulk PCC was

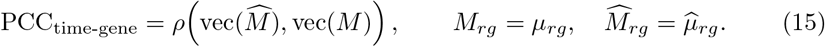

#### Variability and gene co-variation

Gene-variance summaries used all pooled target rows:

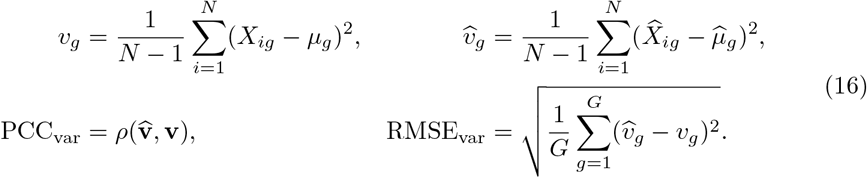

Covariance and variogram calculations used at most 5,000 paired observed/predicted rows per target stage, sampled without replacement using base seed 42 and shared sampling rules across models. Covariance calculations additionally retained a common subset of at most 500 genes. For the resulting observed and predicted sample covariance matrices,

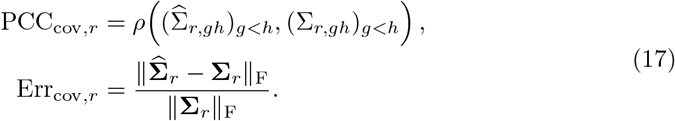

Covariance PCC uses off-diagonal gene pairs, whereas the Frobenius error also includes variances. For sampled rows S*_r_* and a seed-fixed multiset E of 20,000 ordered gene pairs with *g* ≠ *h*, the variogram discrepancy was

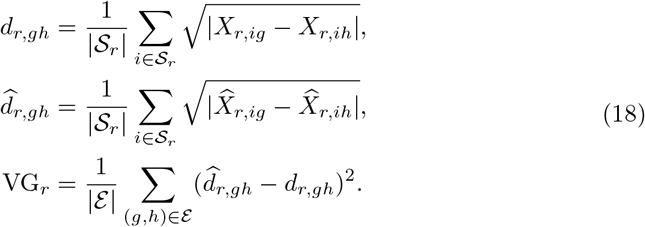

#### Population distributions and expression positivity

Distributional comparisons used at most 2,000 paired rows per stage in an unwhitened PCA space fitted on observed cells and applied unchanged to predictions, retaining at most 30 components. Let **a***_ri_* and **b***_ri_* denote the predicted and observed PCA coordinates, respectively, for *i* = 1*, . . ., n_r_*. For each of 128 seed-fixed unit Gaussian directions, let *a_rℓ,_*_(_*_i_*_)_ and *b_rℓ,_*_(_*_i_*_)_ denote the corresponding sorted projected values. Sliced Wasserstein distance was

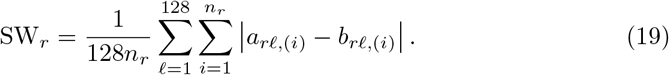

On the same PCA coordinates, the five-kernel U-form squared maximum mean discrepancy was

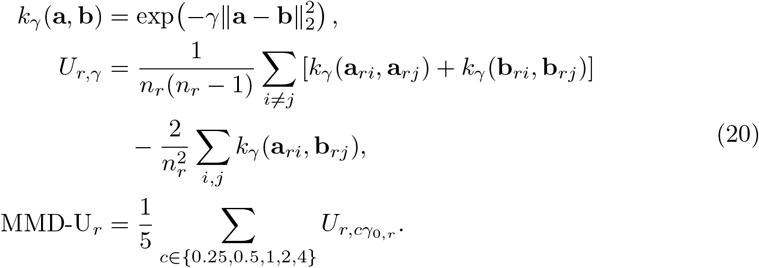

The base parameter *γ*_0_*_,r_* was the inverse median positive squared distance among at most 1,000 observed PCA coordinates. Stagewise population scores were averaged equally across target stages:

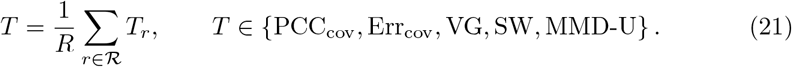

Expression-positivity summaries instead used all pooled target rows:

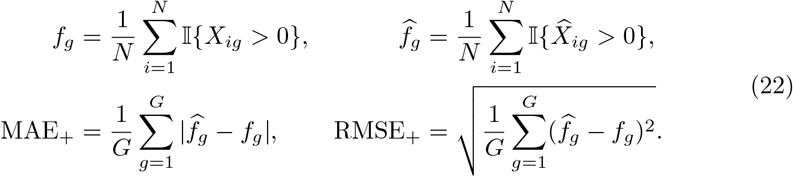

Positivity was defined on the saved expression scale, including centered or shifted representations.

#### Equal-aspect average and radar summary

Within each dataset–panel setting, core endpoints were normalized across BLOOM and all 22 forecasting baselines. For raw endpoint value *b_mj_* of model *m*, define

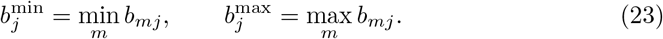

The oriented score was

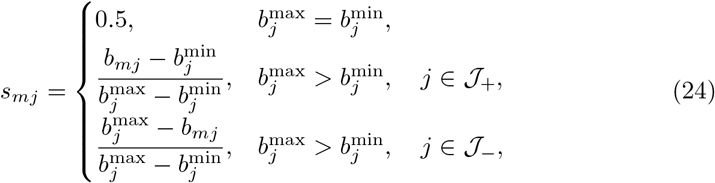

where J_+_ and J_−_ denote higher-is-better and lower-is-better endpoints, respectively. Let *s_mG_*, *s_mE_*, *s_mM_* and *s_mW_* correspond to stage-averaged gene PCC, pooled RMSE, MMD-U and sliced Wasserstein distance. The equal-aspect average assigned equal weight to gene association, expression magnitude and population distributions:

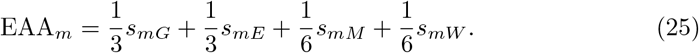

Radar radii used separate endpoint-specific display bounds and clipping, leaving EAA and raw rankings unchanged. The expanded radar displayed an exploratory post-hoc selection of 24 individual endpoints plus EAA, chosen after examining BLOOM’s performance. All 27 candidate endpoints remained in the complete numerical tables, and MMD-U remained in EAA. Scores and rankings were descriptive model comparisons.

### 4.11 Biological expression, cell-state and programme analyses

Biological analyses used observed expression, measured-stage predictions and temporal or perturbation outputs on their native gene panels. Gene means, sample standard deviations and fractions of stored values greater than zero were summarized within developmental stages and supplied cell states. Distribution plots retained cell-level dispersion, while temporal curves summarized population means.

#### Reference expression geometry and cell-state profiles

A PCA reference with at most 30 components was fitted to observed expression and applied unchanged to predictions; UMAP views used the same observed reference with up to 30 neighbours and minimum distance 0.3. Sampled views used stage–state strata and recorded sampling weights for centroids and normalized densities. Density residuals were generated minus observed density on a common grid. Expression-based state maps used a fixed vote among up to five observed neighbours, separately from the state labels supplied to the generator. Marker matrices used full-row state means with a common gene-wise colour transform defined by the mean and standard deviation of observed state means. Cortical dotplots used the same colours and relative positive-expression fractions: a single multiplicative factor per model aligned its mean displayed fraction to the observed mean, preserving within-model ratios.

#### Observed-led gene selection and temporal cascades

Candidate genes came from dataset-derived memberships, supplied programmes and cited developmental evidence. Summary-statistic one-way ANOVA screened observed temporal and stage–state variation. For each gene, the smallest screen *P* value was multiplied by its number of screens and capped at one, followed by Benjamini–Hochberg adjustment across genes. Screening also considered the range of observed group means relative to within-group dispersion. These observed-expression summaries, rather than model prediction errors, guided measured-stage gene selection. For scenarios with only one observed boundary, annotated genes were ranked by generated temporal variation. Cascades ordered genes by their observed peak stage and used the observed temporal mean and standard deviation to scale both observed and generated heatmaps.

The expression and prior screens in Fig. 5f summarized observed temporal variance, state specificity, prior outdegree, prior indegree and membership count in gene programmes curated from the original dataset publications. State specificity was the range of count-weighted state means across time, divided by the pooled observed gene standard deviation. Each component was converted to an average-tie percentile rank over the corresponding 70-, 100- or 135-gene dataset-derived panel before selecting the 25 displayed genes.

#### Programme scoring and gene associations

For programme *p*, represented-gene scores were

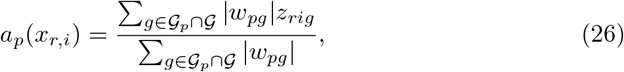

where *w_pg_* = 1 for unweighted memberships. Measured-stage analyses used expression standardized to frozen observed-reference moments when all represented positive-weight members had nonzero reference variance; otherwise the programme used stored expression directly. Temporal-scenario programme scores used stored expression. Scores therefore described the programme members represented in the analysed gene panel. Regulatory association analyses subtracted each gene’s within-stage mean before computing Pearson correlations across the residual expression values. Association error was the absolute difference between the predicted and observed correlations for the same gene pair.

#### Expression-based classifier diagnostics

Candidate cell-state classifiers comprised nearest-centroid classification, class-balanced logistic regression with *C* 0.1, 1, and distance-weighted *k*-nearest-neighbour classification after PCA. Feature standardization and PCA, where applicable, were fitted using training cells. For each evaluation, the classifier was selected by balanced accuracy on observed validation cells and then kept fixed when applied to held-out observed cells and their corresponding generated profiles. Classification performance on held-out observed expression provided a reference for interpreting generated target-label recovery. Confusion-matrix rows denoted supplied evaluation labels and columns denoted classifier assignments; displayed fractions used the full original row support, with unsupported rows left blank. State-fraction diagnostics compared measured annotation fractions, real-expression classifier fractions and generated-expression classifier fractions within the corresponding evaluation windows.

#### Temporal expression summaries and landmarks

Adjacent-stage expression changes were differences between gene-wise population means at consecutive saved stages. Their reported Pearson correlations compared observed and generated changes across the selected gene–stage contrasts. The global expression mean averaged the population-mean vector over genes; variance across that vector described differences among pseudobulk gene values, separately from cell-level variation. Adjacent-time rank persistence was the Spearman correlation between successive population-mean gene vectors. For interpolation, this rank diagnostic compared generated query rows after excluding observed anchors and identified comparisons spanning an omitted anchor.

For a gene with population means *m_k_* at ordered coordinates *t_k_*, the maximum was the first coordinate attaining max*_k_ m_k_*. The half-range threshold and endpoint direction were

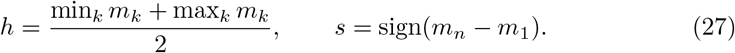

The first crossing satisfied *s*(*m_k_*_−1_ − *h*) *<* 0 and *s*(*m_k_ h*) 0; its coordinate was obtained by linear interpolation between the two bracketing values. A crossing was reported only if the profile began below the threshold in that direction, *s*(*m*_1_ − *h*) *<* 0. Boundary maxima were identified separately, and the descriptor analysis distinguished observed-anchor maxima from generated-query maxima. Each component retained its recorded stage, age or reporting coordinate.

#### Summaries of single-gene input responses

Perturbation views used edited-minus-baseline expression differences from matched model calls. Downstream response magnitude was the root mean square over non-edited genes; marker-set summaries also excluded the edited gene. Signed heatmaps displayed population-mean responses on fixed 25-gene panels. In association diagrams, each responding gene was coloured at its own stage of maximum absolute response.

## Competing interests

The authors declare no competing interests.

## Data availability

This study analyzed three publicly available mouse single-cell transcriptomic datasets. The mouse gastrula-to-birth atlas [8] is available from the Gene Expression Omnibus under accession numbers GSE186069 and GSE228590, with processed data and annotations also accessible at https://omg.gs.washington.edu/. The developing mouse visual-cortex atlas [9] is available through the Allen Brain Cell Atlas. The STED-EC atlas [10] is available from the National Genomics Data Center under accession OMIX007369. No new sequencing data were generated in this study.

## Code availability

The BLOOM source code and trained model checkpoints will be made publicly available at https://github.com/tracy666/BLOOM.

## Author contribution

LY and YH conceived, designed, and supervised the study. ZL developed and implemented the BLOOM framework, conducted the experiments, and performed data analyses. ZL, SL, BL, LY, and YH wrote the manuscript, with critical input from all authors. All authors contributed to discussions and approved the final manuscript.

## Notes

### Competing Interest Statement

The authors have declared no competing interest.

